# An FcRn-targeted mucosal vaccine against SARS-CoV-2 infection and transmission

**DOI:** 10.1101/2022.11.23.517678

**Authors:** Weizhong Li, Tao Wang, Arunraj M. Rajendrakumar, Gyanada Acharya, Zizhen Miao, Berin P. Varghese, Hailiang Yu, Bibek Dhakal, Tanya LeRoith, Wenbin Tuo, Xiaoping Zhu

**Affiliations:** Division of Immunology, VA-MD College of Veterinary Medicine, *Maryland Pathogen Research Institute, University of Maryland, College Park, MD 20742; Animal Parasitic Diseases Laboratory, ARS, United States Department of Agriculture, Beltsville, MD 20705; Department of Biomedical Sciences and Pathobiology, Virginia-Maryland Regional College of Veterinary Medicine, Virginia Tech University, Blacksburg, VA, USA

**Keywords:** COVID-19, SARS-CoV-2, vaccine, mucosal, nasal, FcRn, mouse, hamster

## Abstract

SARS-CoV-2 and its variants cause COVID-19, which is primarily transmitted through droplets and airborne aerosols. To prevent viral infection and reduce viral spread, vaccine strategies must elicit protective immunity in the airways. FcRn transfers IgG across epithelial barriers; we explore FcRn-mediated respiratory delivery of SARS-CoV-2 spike (S). A monomeric IgG Fc was fused to a stabilized S protein; the resulting S-Fc bound to S-specific antibodies (Ab) and FcRn. A significant increase in Ab responses was observed following the intranasal immunization of mice with S-Fc formulated in CpG as compared to the immunization with S alone or PBS. Furthermore, we intranasally immunize adult or aged mice and hamsters with S-Fc. A significant reduction of virus replication in nasal turbinate, lung, and brain was observed following nasal challenges with SARS-CoV-2, including Delta and Omicron variants. Intranasal immunization also significantly reduced viral transmission between immunized and naive hamsters. Protection was mediated by nasal IgA, serum-neutralizing Abs, tissue-resident memory T cells, and bone marrow S-specific plasma cells. Hence FcRn delivers an S-Fc antigen effectively into the airway and induces protection against SARS-CoV-2 infection and transmission. Based on these findings, FcRn-targeted non-invasive respiratory immunizations are superior strategies for preventing highly contagious respiratory viruses from spreading.

## Introduction

SARS-CoV-2, a virus causing the COVID-19 pandemic, is highly infectious and circulates rapidly worldwide, and mutates constantly. The SARS-CoV-2 variants include Alpha (B.1.1.7), Beta (B.1.351), Gamma (P.1), Delta (B.1.617.2), and Omicron (B.1.1.529). These variants evade the immunity induced by vaccinations and infections. Through the receptor-binding domain (RBD) of its spike (S) protein, SARS-CoV-2 binds to the angiotensin-converting enzyme 2 (ACE2). The S protein undergoes a structural change upon binding and is cleaved by host proteases, such as the transmembrane serine protease 2 (TMPRSS2), allowing it to fuse with the cellular membrane for host cell entry^1^. Because of its vital role in mediating receptor binding and infection initiation, the S protein is the primary target for developing vaccines^2^.

SARS-CoV-2 can be shed from individuals with asymptomatic infections and spread predominantly through droplets and airborne aerosols^3^. The virus first enters the nose or mouth and replicates within epithelial cells of the nasopharynx, causing an upper respiratory infection. Hence, the nasal mucosa and nasopharynx are the primary sites of exposure to SARS-CoV-2 before dissemination to the lungs and other tissues/organs^4^. The currently authorized intramuscular vaccines can effectively prevent severe diseases and deaths caused by COVID-19. However, they are unable to effectively elicit protective mucosal immunity in the upper respiratory tract^5–7^. This shortcoming allows opportunistic breakthrough infections in those who received vaccinations^8^. Hence, the SARS-CoV-2 can linger in the nasal mucosa even after clearing infection in the lungs in vaccinated individuals. The emergence of the SARS-CoV-2 variants, especially the Omicron, exacerbates the situation. This ongoing evolution of SARS-CoV-2 necessitates a safe and protective mucosal vaccine to block the viral entry and reduce or eliminate the viral spread, thus, prevent lung and systemic infection and breakthrough infection. Nasal-spray vaccines can elicit local secretory IgA antibodies and resident T and B cell responses in the upper respiratory tract and the lungs^6, 9, 10^.

Epithelial cells lining the respiratory tract form a mucosal barrier. The neonatal Fc receptor (FcRn) binds to the Fc portion of IgG and mediates the transfer of IgG across the epithelial cells, a function essential to IgG distribution and homeostasis^11–14^. Typically, FcRn shows a pH-dependent IgG binding, with a preference to bind IgG at acidic pH (6.0 – 6.5)^14^. Generally, the FcRn-IgG on the cell surface or in the endosome under acidic conditions goes through a non-degradative vesicular transport pathway within epithelial cells. Consequently, FcRn transports its bound IgG across the mucosal barrier and then releases it into the lumen or submucosa upon exposure to physiological pH^14^. Through binding, FcRn also extends the half-life of IgG by reducing lysosomal degradation in cells, such as endothelial cells^15^.

The vaccine must be administered locally in the respiratory tract to establish respiratory immunity with resident memory of T and B cells in the lungs^16, 17^. In this study, we determined the ability of FcRn to deliver an intranasally-administered SARS-CoV-2 S antigen and induce protective mucosal and systemic immunity to SARS-CoV-2 infection. We defined protective immune responses and mechanisms relevant to this nasal vaccination in the mouse and hamster models. Our results show that FcRn-mediated nasal delivery of a prefusion-stabilized SARS-CoV-2 S antigen induces secretory IgA Abs in the nasal lavage and high levels of long-lasting Ab and T-cell responses. We found that our nasal vaccine conferred durable protection against SARS-CoV-2 infection and transmission.

## Results

### Expression and characterization of SARS-CoV-2 S-Fc fusion proteins

To target antigen to FcRn, we expressed a monomeric human IgG1 Fc fused to a prefusion-stabilized, soluble form of S, which contained an R685A mutation at the furin cleavage site, an R816A mutation at S2’ cleavage site, and 2P conversions (Fig. 1a) along with the T4 fibritin trimerization domain^18^. We also removed the C1q binding site in IgG1 Fc.

**Figure 1.**
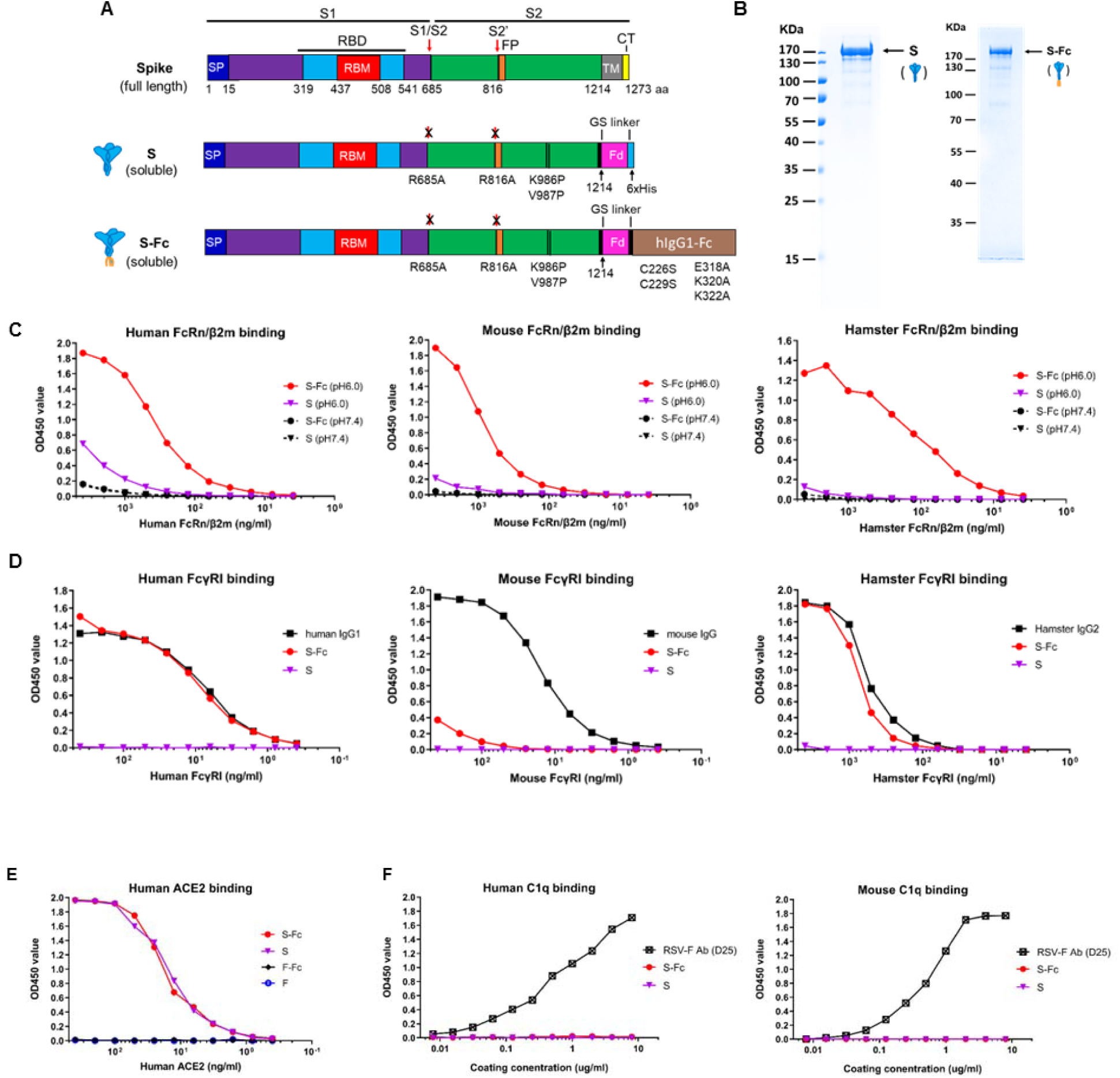
Expression and characterization of the prefusion-stabilized Spike-IgG Fc (S-Fc) fusion proteins. **A.** Schematic illustration of the full-length protein sequence of SARS-CoV-2 Spike (*top*). SP: signal peptide; RBD: receptor binding domain; TM: transmembrane domain; CT: cytoplasmic tail. Graphic design of the fusion of SARS-CoV-2 Spike with the T4 fibritin foldon domain (Fd) to create a soluble, prefusion-stabilized, and trimeric Spike protein (*middle*). Mutations were made in SARS-CoV-2 (strain USA/WA1/2020) spike by replacing Arg 685 and Arg 816, respectively, with an Ala residue to remove the cleavage site, and replacing Lys 986 and Val 987, respectively, with a Pro residue to create a prefusion-stabilized form. Diagram demonstration of the fusion of a SARS-CoV-2 Spike with the T4 fibritin foldon domain (Fd) and human Fcγ1 to create a prefusion-stabilized and trimeric S-Fc fusion protein (*bottom*). Mutations were also made in the Fcγ1 fragment by replacing Cys 226 and Cys 229, respectively, with a Ser residue to abolish Fc dimerization, and replacing Glu 318 Lys 320 Lys 322 with Ala residues to remove the complement C1q binding site. **B.** The S and S-Fc fusion proteins were purified from the stable CHO cell lines. The soluble S (*left*) and S-Fc (*right*) proteins were purified by anti-His and Protein A affinity chromatography respectively and subjected to SDS-PAGE gel electrophoresis under reducing conditions and visualized with Coomassie blue staining. The molecular weight in kDa is marked in the left margin. **C+D+E+F**. Test of the S-Fc binding to human, mouse, or hamster FcRn/β2m; human, mouse or hamster FcγRI; human ACE2; and human or mouse C1q. The specific binding was determined by the ELISA. The purified S protein was used as a positive control for ACE2 binding and negative control for FcRn/β2m or FcγRI binding. Respiratory syncytial virus (RSV) protein F alone or Fc-fused F proteins were used as a negative control. Mouse IgG, human IgG1, hamster IgG2, and a mAb (D25) against RSV F protein were used as the positive control, respectively.

We showed that the soluble S or S-Fc protein was secreted from the stable CHO cells (Fig. 1b). Since IgG only binds to FcRn at acidic pH^14^, we determined if the S-Fc portion binds to FcRn at either pH 6.0 or 7.4. As shown in Fig. 1c, S-Fc interaction with both human (Fig. 1c) and mouse (Fig. 1c) FcRn/β2m was detected at pH 6.0 condition. Since hamster FcRn/β2m is not available, we produced a biotinylated hamster FcRn/β2m (Extended Fig. 1d). We found the S-Fc bound to hamster FcRn/β2m similar to human or mouse FcRn/β2m did (Fig. 1c). However, the binding of the S to human, mouse, or hamster FcRn/β2m protein was barely detectable (Fig. 1c). To further show whether the S-Fc binds to FcγRI (CD64), we also produced biotinylated hamster FcγRI (Extended Fig. 1e). As expected, the S protein did not bind to mouse, human, or hamster FcγRI (Fig. 1d). As shown in Fig. 1d, the S-Fc, human IgG1, or hamster IgG2 could bind human or hamster FcγRI similarly in an ELISA assay. However, the S-Fc did not bind mouse FcγRI with high affinity (Fig. 1d), although mouse IgG was shown to bind mouse FcγRI strongly. This result could be explained by the fact that human IgG1 does not interact with mouse FcγRI with high affinity^19^.

We next determined if the S portion of the S-Fc binds to human ACE-2. In an ELISA assay, the S-Fc and S protein bound human ACE-2 similarly (Fig. 1e), indicating the Fc fusion doesn’t affect the conformation of S in the S-Fc protein. As a negative control, RSV F protein with or without Fc-fusion did not bind to human ACE-2 (Fig. 1e). In contrast to an RSV-F specific mAb (D25), the S-Fc could not bind to human or mouse C1q protein (Fig. 1f) in *vitro*. We further wanted to know if the S portion of the S-Fc interacts with convalescent serum Abs compared to the S alone. Normal human sera from the different healthy donors were used as a negative control. Indeed, the sera from convalescent COVID-19 patients (Extended Fig. 2a) were able to equivalently recognize both purified S-Fc (*solid lines*) and S (*dashed lines*) protein, showing varying binding efficiencies. Most importantly, S-Fc and S showed the same levels of binding for each tested serum sample, indicating Fc-fusion with S didn’t alter its conformation. Normal sera (Extended Fig. 2b) from healthy individuals showed no binding activity. The results were further confirmed by the interactions of both S-Fc and S with various spike-specific mAbs (Extended Fig. 2c, d). Together, the S portion of the S-Fc protein maintains the correct conformational structure allowing for binding to the ACE-2 and S-specific Abs and also possesses the function to engage with FcRn and human or hamster FcγR1.

### FcRn-dependent respiratory immunization significantly enhances Spike-specific immune responses

We tested whether FcRn-dependent transport augments the immune responses to S protein. C57BL/6 mice were i.n. immunized with 10 μg of S-Fc, S protein (equal molar amount), or PBS in 10 μg CpG adjuvant, the mice were boosted after two weeks (Fig. 2a). FcRn-knockout (KO) mice were used as a control to test the FcRn-mediated immunity enhancement. Using S protein alone allowed us to evaluate FcRn-independent effects *in vivo* and determine the magnitude of enhanced immune responses conferred by targeting the S-Fc to FcRn. As shown in Fig. 2, significantly higher titers of total serum IgG (Fig. 2b, p<0.0001) were detected in the S-Fc immunized mice compared with the S-Fc-immunized FcRn KO, the S-immunized or PBS-treated mice. Moreover, sera from the S-Fc immunized mice exhibited much stronger neutralizing activity relative to the control groups (Fig. 2c, *p<0.01-0.0001*). Likewise, the S-Fc induced higher levels of IgG (Fig. 2d, e) and IgA (Fig. 2f, g) Abs in nasal washes (Fig. 2d, f) and bronchoalveolar lavage fluids (BAL) (Fig. 2e, g) than those of the mice immunized by the S alone, PBS or the FcRn KO mice immunized by the S-Fc.

**Figure 2.**
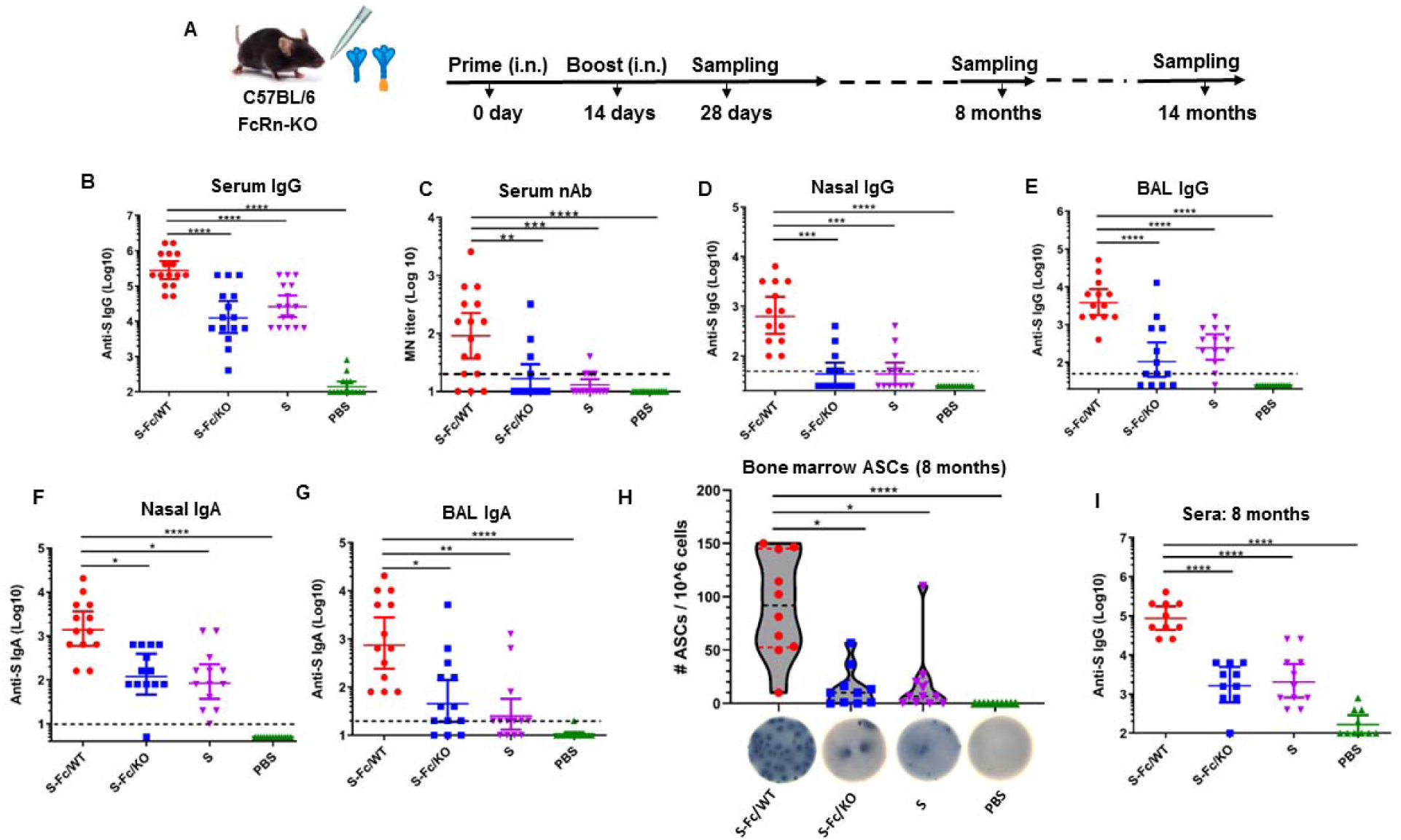
FcRn-mediated respiratory immunization induces SARS-CoV-2 S-specific antibody immune responses. **A.** Ten μg of S-Fc, S (with the equivalent molar number), or PBS in combination with 10 μg of CpG was i.n. administered into 6-8-week-old wild-type (WT) C57BL/6, or FcRn knockout (KO) mice. Mice were boosted 14 days after the primary immunization. Sampling was performed at the indicated time points. **B.** Anti-SARS-CoV-2 S-specific IgG Ab titers in sera. The S-specific Ab titers were measured by coating the plates with S protein in ELISA. The IgG titers were measured in 14-16 representative mouse sera per group. The data represent a geometric mean with 95% CI. **C.** Test of neutralizing Ab activity in sera of the immunized animals. Sera from 14 to 16 mice per group were heat-inactivated and serially diluted two-fold. SARS-CoV-2 (100 TCID_50_) was added and incubated at 37°C for 1 hr. The mixture was added to Vero-E6 cells and incubated at 37°C for 96 hrs, the neutralization Ab titers were determined and are expressed as the reciprocal of the highest dilution preventing the appearance of the cytopathic effect (CPE). **D, E, F & G**. Anti-S-specific Ab titers in nasal washings (**D**, IgG; **F**, IgA), and BAL (**E**, IgG; **G**, IgA). The data represent a geometric mean with 95% CI. **H**. S-specific Ab-secreting cells in the bone marrow. Bone marrow cells isolated 8 months after the boost was placed on S-coated plates and quantified by ELISpot analysis of IgG-secreting plasma cells. Data were pooled from two separate experiments for at least 10 immunized mice in each group. The graphs were plotted based on each experiment’s average spot number from 6 wells. **I**. Anti-SARS-CoV-2 S-specific IgG Ab titers in sera 8 months after the booster. The data represent a geometric mean with 95% CI. One-way ANOVA with Dunnett’s multiple comparison tests was used. A nonparametric test was used if the data were not normally distributed. Asterisks in the figures: *, P < 0.05; **, P < 0.01; ***, P<0.001; ****, P<0.0001.

Activated B cells can differentiate into plasma cells that secrete antibodies at a high rate and reside in niches in the bone marrow. To determine whether antigen targeting to FcRn also elicited plasma cells that secreted S-specific Abs, the number of IgG-secreting plasma cells in the bone marrow was measured 8 months after the boost by ELISpot. High numbers of S-specific IgG-secreting cells were present in the bone marrow of mice immunized with S-Fc compared with other groups (Fig. 2h, *p<0.05*). To show whether increased IgG-secreting plasma cells correspond to a rise in Ab production and maintenance, IgG Abs in the sera were measured 8 months after the boost. High titers of S-specific IgG Abs were maintained in mice immunized with the S-Fc, but not the S alone (Fig. 2i). Also, a significant level of S-specific IgA or IgG was present in the nasal washes or BAL of mice immunized with S-Fc compared with other groups 14 months later (Extended Fig. 3a-d). Immunization with the S-Fc was more effective than immunization with S alone, indicating that S-specific Abs persisted much longer after FcRn-targeted mucosal immunization. Overall, our data demonstrate that an Fc-fused, soluble, prefusion-stabilized S protein delivered through FcRn is much more potent in triggering S– specific Ab responses.

### Intranasal vaccination by the S-Fc leads to increased protection against infection by the ancestral SARS-CoV-2 in K18-hACE2 transgenic mice

Human ACE2 transgenic mice are highly susceptible to SARS-CoV-2 intranasal challenges when high virus doses are used^20^. In the study, hACE2 mice were i.n. immunized with 10 μg of S-Fc or PBS in 10 μg CpG and boosted in a 2-week interval (Fig. 3a). We confirmed that significantly higher titers of serum IgG (Fig. 3b, P<0.0001) and neutralizing antibodies (nAbs) (Fig. 3c, P<0.01) were detected in the hACE2 mice i.n. immunized with the S-Fc when compared with PBS-treated mice. To test whether the immune responses elicited by the intranasal (i.n.) vaccination with the S-Fc provide protection, we i.n. challenged all immunized mice with a lethal dose (2.5 X 10^4^ TCID_50_) of ancestral SARS-CoV-2 virus 2-3 weeks after the boost (Fig. 3a). Mice were monitored and weighed daily for 14 days. All mice in the control groups exhibited rapid weight loss following the challenge, either succumbing to infection within 8 days post-infection (dpi) or subjecting to euthanasia. In contrast, the S-Fc-immunized mice did not experience significant body-weight loss (Fig. 3d) and they had full protection with significantly higher survival rates (100%) than those of the PBS control group (Fig. 3e). As expected, all the PBS-treated mice had high virus loads in the lung at 5 dpi. In contrast, a significant reduction of virus load in the nasal turbinate (P<0.0001), lungs (P<0.0001), and brain tissue (P<0.0001) were seen in the S-Fc immunized mice when compared to the control animals (Fig. 3f). Interestingly, brain tissues exhibited the highest levels of virus load in the PBS control mice at 5 dpi. Therefore, the S-Fc-immunized hACE2 mice essentially contained viral replication in tissues/organs of the viral entry and prevented the viral spreading to other tissues/organs.

**Figure 3.**
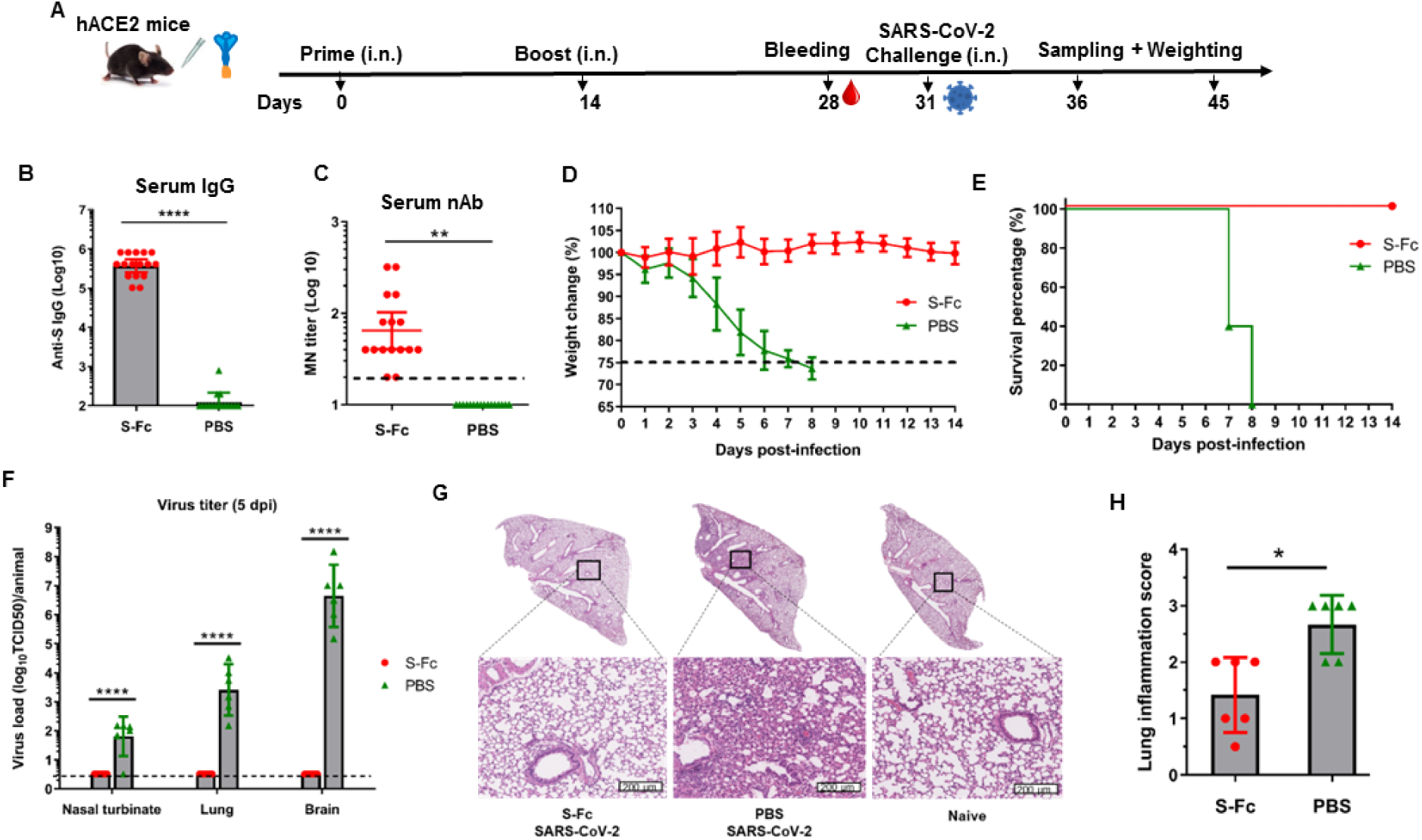
Intranasal immunization by the S-Fc induces protective immunity to intranasal (i.n.) challenge with ancestral SARS-CoV-2 virus. **A.** 10 μg of S-Fc or PBS in combination with 10 μg of CpG was i.n. administered into 8-week-old hACE-2 mice (n=15-16). Mice were boosted 14 days after primary immunization. Five to six mice in each group are euthanized at 5 dpi for sampling and virus titration. The remaining 10 mice in each group are subjected to the survival analysis. **B & C**. Serum anti-SARS-CoV-2 S-specific IgG Ab titers (**B**) and neutralizing Ab (**C**) in hACE2 mice. The S-specific IgG Ab titers were measured by coating the plates with S protein in ELISA; the neutralizing Ab activity in the immunized sera was determined by the micro-neutralization test. The data represent a geometric mean with 95% CI. **D.** Body-weight changes following the SARS-CoV-2 challenge. Seventeen days after the boost, groups of 10 mice (S-Fc group, n=10; PBS group, n=10) were i.n. challenged with the SARS-CoV-2 virus (2.5 X 10^4^ TCID_50_) and weighed daily for 14 days. Mice were humanely euthanized at the end of the experiment or when a humane endpoint was reached. **E.** Survival following virus challenge. The percentage of mice protected after the challenge was shown by the Kaplan-Meier survival curve. **F.** Viral titers in the nasal turbinate, lung, and brain 5 days after the challenge. The virus titers in the samples of the immunized and control mice (n=5-6) were determined. The presence of live virus was determined by CPE in Vero E6 cells cultured for 4 days. The viral titers were shown as TCID_50_ from each animal sample. **G.** Histopathology of the lungs from the challenged and naïve mice. Lungs were collected 5-days post-challenge. The lungs from uninfected mice were included as normal control (n=6). The lung sections were stained with Hematoxylin-Eosin (H & E) to determine the intensity of inflammation. The representative slides are shown. All scale bars represent 200 µm. **H.** The inflammatory responses of each lung section were scored in a blind manner. The student t-test was used for the statistical assay.

To further show protection, lungs were collected at 5 dpi for histopathological analysis. The lungs of uninfected mice were used as normal control (Fig. 3g). Prior to the virus challenge, no apparent alterations were observed in the lung structure and histology of the S-Fc immunized mice and normal mice, suggesting that the S-Fc did not induce inflammation. In contrast, we found focal inflammatory cell infiltration, pneumonia, peribronchiolitis, and perivasculitis in the lungs of PBS control mice after virus infection (Fig. 3g). The alveolitis was not observed. Hence, the mice immunized with S-Fc had a significantly lower inflammation score of the lungs compared with those of mice in the PBS group (*Fig. 3h,* P<0.05). We also found that i.n. immunization with the S-Fc protected the aged hACE2 mice from lethal SARS-CoV-2 infection (Extended Fig. 4a-f; Supplementary Information A) and elicited durable protection in hACE2 mice (Extended Fig. 4g-j; Supplementary Information B). These findings demonstrate that FcRn-mediated delivery of the S-Fc confers significant protection against lethal SARS-CoV-2 challenge, resulting in decreased mortality, viral replication, and pulmonary inflammation in a hACE2 mouse model.

### Intranasal vaccination with the S-Fc protein leads to protection against SARS-CoV-2 Delta and Omicron variants

The SARS-CoV-2 is rapidly evolving via mutagenesis, which significantly impacts transmissibility, morbidity, reinfection, and mortality and lengthens the pandemic. Since the S portion of the S-Fc is derived from ancestral SARS-CoV-2, we were interested in determining the effectiveness and neutralizing activity elicited by the S-Fc vaccine against both Delta and Omicron variants. First, we i.n. immunized hACE2 mice twice with 10 μg of S-Fc adjuvanted in 10 μg CpG (Fig. 4a) and tested the protection against the SARS-CoV-2 Delta strain. The majority of the immunized mice developed nAbs against the Delta strain after the boost (Extended Fig. 3e, p<0.01). To show protection, we challenged all immunized mice with a lethal dose (2.5 x 10^4^ TCID_50_) of the Delta strain 17 days after the boost. All mice in the control group experienced rapid weight loss (Fig. 4b), labor breathing, and ataxia, and finally died from the viral infection, or were euthanized for humanity. In contrast, the S-Fc-immunized hACE2 mice did not show significant body-weight loss (Fig. 4b) or clinical signs. The majority (83.3%) of the S-Fc-immunized mice survived, which was significantly higher than the survival rates of the PBS control groups, where all mice died from Delta virus infection (Fig. 4c). Further, we measured viral replication in the nasal turbinate, lung, and brain tissues 6 DPI (Fig. 4d). We were able to detect live Delta virus in the nasal turbinate, lung, and brain tissues of the PBS control mice, but failed to find any live virus in the nasal turbinate and lung tissues of the S-Fc-immunized animals (Fig. 4d). For brain samples, only one mouse in the S-Fc-immunized group showed the reduced virus load, while the others had no virus detected. Additionally, no prominent inflammation was observed in the lungs of the S-Fc immunized mice. In contrast, focal perivascular and peribronchial inflammation and thickened alveolar septa were found in the lung of PBS control mice (Fig. 4e). The mice immunized with S-Fc had a significantly lower inflammatory score compared with those of mice in the challenged animals in PBS control group (Extended Fig. 3f, P<0.01).

**Figure 4.**
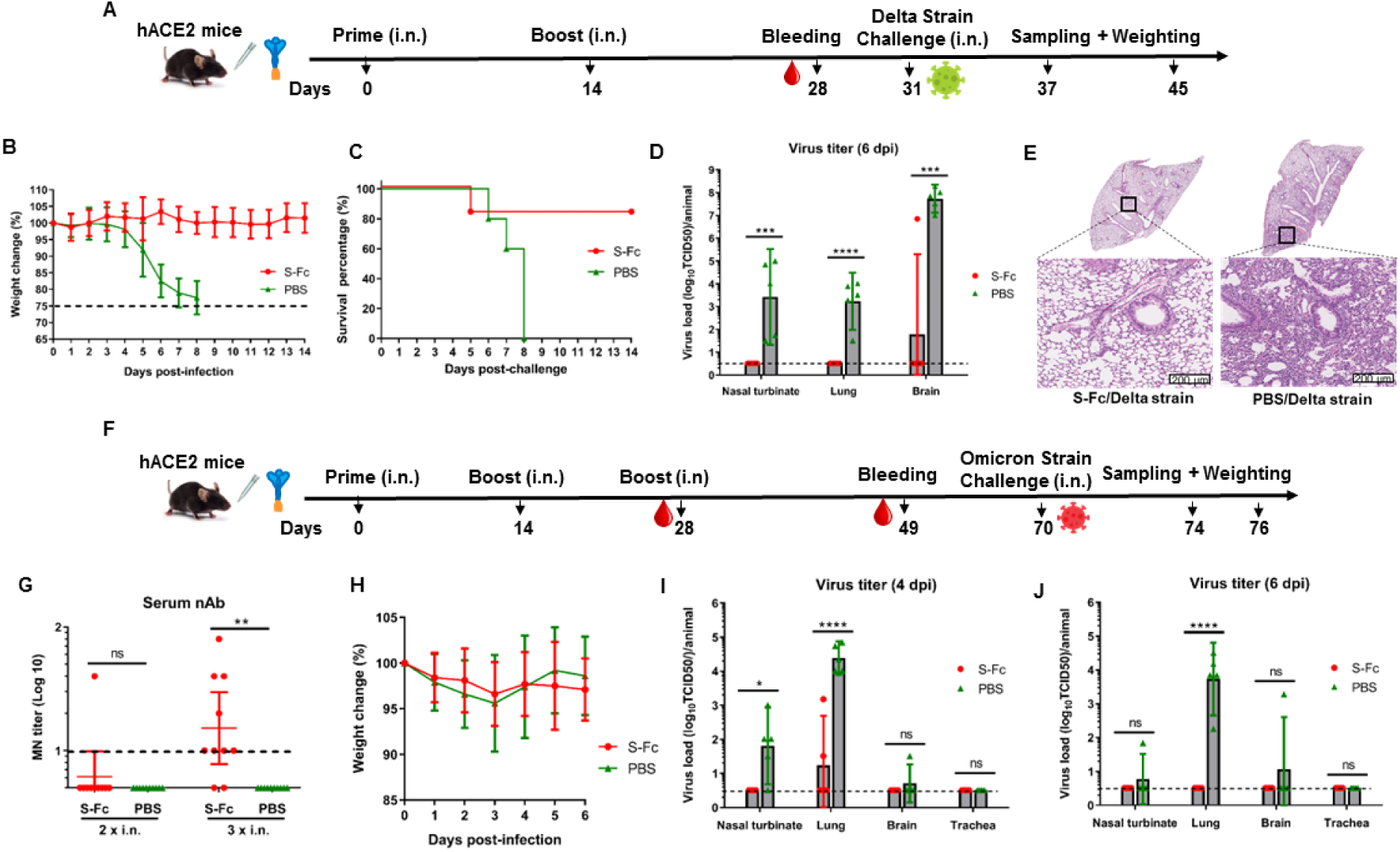
Intranasal immunization by the S-Fc induces protective immunity to intranasal challenge with SARS-CoV-2 Delta or Omicron variants. **A.** 10 μg of S-Fc or PBS in combination with 10 μg of CpG was i.n. administered into 8-week-old hACE-2 mice (n=10-11). Mice were boosted 14 days after the primary immunization. Five mice in each group are euthanized at 5 dpi for sampling and virus titration. The remaining 5-6 mice in each group are subjected to the survival analysis. **B.** Body-weight changes following the SARS-CoV-2 Delta variant challenge. 17 days after the boost, groups of mice (S-Fc group, n=6; PBS group, n=5) were i.n. challenged with SARS-CoV-2 Delta strain (2.5 X 10^4^ TCID_50_) and weighed daily for 14 days. Mice were euthanized at the end of the experiment or when a humane endpoint was reached. **C.** Survival following virus challenge. The percentage of hACE-2 mice protected after the virus challenge was shown by the Kaplan-Meier survival curve. **D.** Viral titers in the nasal turbinate, lung, and brain 6 days after Delta strain challenge. The live virus titers in the tissues of the S-Fc immunized and PBS control mice (n=5) were determined. Supernatants of the nasal turbinate, lung, and brain homogenates were added to Vero-E6 cells and incubated for 4 days. The viral titers were shown as TCID_50_ from each animal sample. **E.** Histopathology of the lungs from the challenged hACE-2 mice. Lungs were collected 6-day post-challenge. The lung sections were stained with Hematoxylin-Eosin (H & E) to determine the level of inflammation. The representative slides were shown. All scale bars represent 200 µm. **F.** Ten μg of S-Fc or PBS in combination with 10 μg of CpG was i.n. administered into 8-weeks old hACE-2 mice (n=10). Mice were boosted 14 and 28 days after primary immunization, respectively. Mice were i.n. challenged with the Omicron variant 42 days after the final boost. Five mice from each group are euthanized at 4 and 6 dpi for sampling and titrating the virus. **G.** Neutralizing Ab (nAb) in the immunized mouse sera. Two weeks after the first boost or three weeks after the second boost, sera sampled from 10 mice per group were heat-inactivated and serially diluted two-fold in PBS. SARS-CoV-2 Omicron A.1 strain (100 TCID_50_) was added and incubated at 37°C for 1 hr. The mixture was added to VAT cells; after incubation at 37°C for 96 hr, the nAb titers were determined and expressed as the reciprocal of the highest dilution preventing the appearance of the CPE **H.** Body-weight changes following the Omicron A.1 challenge. Forty-two days after the boost, groups of 10 mice (S-Fc group, n=10, PBS group, n=10) were i.n. challenged with Omicron A.1 strain (1 X 10^6^ TCID_50_) and weighed daily for 4 or 6 days. Half mice in each group (n=5) were euthanized at 4 dpi and another half mice (n=5 for each group) were euthanized at 6 dpi. **I+J.** Viral titers in the nasal turbinate, lung, and brain 4 or 6 days after Omicron A.1 virus challenge. The virus titers in the tissues of the immunized and control mice (n=5) were determined. Supernatants of the nasal turbinate, lung, and brain homogenates were added to VAT cells and incubated for 4 days. The viral titers were shown as TCID_50_ from each animal sample. The student *t*-test was used for the statistical assay.

Interestingly, three i.n. immunizations were necessary to produce the Omicron-specific nAbs in the sera of the majority of mice (Fig. 4f, g). Very likely, the Omicron strain possesses abundant mutations in its S protein, implicating that Omicron can escape nAbs elicited by the ancestral S protein^21^. Noticeably, when the S-Fc immunized mice were challenged with a dose (1 X 10^6^ TCID_50_) of the Omicron A.1 strain after 2nd boost, there was no difference in visual clinical symptoms and body-weight loss between the S-Fc-immunized and the PBS control mice throughout the experiments (Fig. 4h), which can be attributed to the low virulence of Omicron strain in mice. We did not find any live Omicron virus in nasal turbinate and brain tissues (Fig. 4i, j) in all S-Fc-immunized mice at 4 and 6 dpi. For lung tissues, the live Omicron virus was only identified in 2 mice immunized with S-Fc at 4 dpi at low levels (around 1030 and 22 folds of virus reduction compared to the average virus titer in PBS groups) but not 6 dpi, indicating the limited virus replication in these mice. Furthermore, we found a dramatically decreased inflammation in the lung of the S-Fc immunized mice compared with the PBS control mice who had apparent focal perivascular and peribronchial inflammation. Overall, in contrast to the ancestral and Delta strains, infection of the Omicron A.1 in mice did not cause body-weight loss and mortality. However, the S-Fc nasal immunization remarkably attenuated replication of the Omicron variant in the respiratory tract and the brain, and resulted in substantially reduced lung inflammation in the hACE2 mice.

### Intranasal immunization with the S-Fc protein induces local immune responses and abrogates SARS-CoV-2 viral replications in the respiratory tract

Because SARS-CoV-2 initiates its infection in the upper respiratory tract^3^, it is critical for a nasal vaccine to elicit anti-viral IgA Abs in the nasal secretions and BAL of the lung. First, to determine the ability of the respiratory immunization by the S-Fc to induce local humoral immune responses. We examined S-specific Abs in mucosal secretions, which were compared to those of mice that received intramuscular (i.m.) immunization with the same amount of the S-Fc and CpG (Fig. 5a). The nasal washes and BAL were collected 14 days following the boost and analyzed for S-specific IgA and IgG by ELISA. As shown in Fig. 5, the levels of S-specific IgA Abs significantly increased in the nasal washes and BAL (Fig. 5b, c) in the i.n. immunized mice. In contrast, the mice that were immunized with the S-Fc via the i.m. route showed much lower levels of S-specific IgA in both the nasal washes and BAL (Fig. 5b, p <0.0001). The mice that were i.m. immunized with the S-Fc had a higher level of S-specific IgG in the nasal washes than that of the i.n. immunized mice (Fig. 5d, p<0.01). However, both i.n. and i.m. S-Fc-immunized mice exhibited similar levels of IgG Ab in the BAL (Fig. 5c). The mice that were i.m. immunized with the S-Fc also developed higher levels of IgG Ab in the sera than those of the i.n. immunized mice (Fig. 5d, p<*0.01*); this may reflect the full deposit of the S-Fc protein in the tissue by the needle injection, compared to i.n. immunization which usually resulted in a lower-than-desired dose.

**Figure 5.**
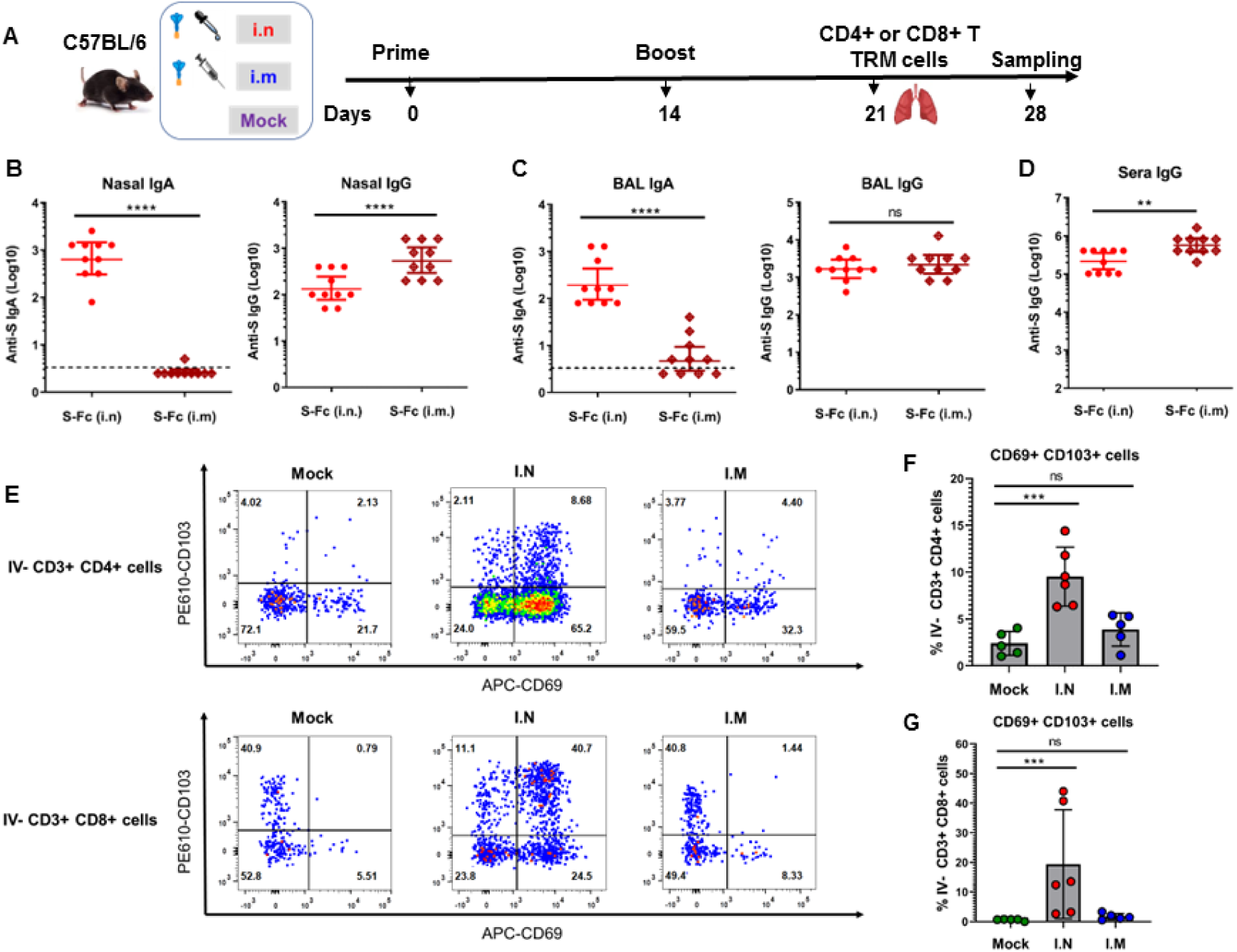
FcRn-mediated intranasal vaccination significantly induces S-specific local immune responses in the respiratory tract. **A.** Ten μg of S-Fc or PBS in combination with 10 μg of CpG was i.n. administered into 6-8 week old mice (n=10). Mice were boosted 14 days after primary immunization. An additional group of mice (n=10) that were intramuscularly (i.m.) immunized in 10 μg S-Fc with 10 μg of CpG was included as an i.m. route control. Tissues were collected at the indicated time points. **B+C+D**. Anti-SARS-CoV-2 S-specific IgA or IgG Ab titers in nasal washings (**A**), BAL (**B**), and serum (**C**) after the boost. SARS-CoV-2 S-specific Abs were measured by ELISA 14 days after the boost in 10 representative mouse samples per group. The data represent a geometric mean with 95% CI. **E+F+G.** Tissue-resident memory (TRM) T cells in mouse lungs. The IV-CD3^+^CD4^+^CD69^+^ CD103^+^ (*top panel*) or IV-CD3^+^CD8^+^CD69^+^CD103^+^ (*bottom panel*) TRM T cells in the lungs were assessed 7 days after the boost by FACS. Flow cytometry plots are the representative results from 5-6 individual samples per group. Numbers in the top right quadrants (**E**) or the graph (**F+G**) represent the percentage of TRM CD4^+^ or CD8^+^ T lymphocytes following i.n. or i.m. immunizations. The student t-test was used for the statistical analysis of data from experiments B-D, while one-way ANOVA was used for analyzing the results of F and G. Nonparametric test was used if the distribution of data was not normal.

Tissue-resident memory (TRM) T cells are found in the nasal cavity and lungs during SARS-CoV-2 infection, which is essential to limit disease severity and viral replication^22^. Thus, we next determined whether intranasal delivery of the S-Fc protein can induce TRM T cells in the lungs. The TRM T cells in the lungs were assessed 7 days after the boost by FACS (Fig. 5e and Extended Fig. 5). Compared to the PBS control, we detected a notably higher percentage of CD4^+^CD69^+^CD103^+^ TRM cells (Fig. 5f) and CD8^+^CD69^+^CD103^+^ TRM cells (Fig. 5g) in the lungs of the i.n. immunized, but not in the i.m. immunized mice with the S-Fc. Together, these data suggest that the intranasal, but not intramuscular, immunization with the S-Fc protein induces CD4^+^ and CD8^+^ TRM cells in the lung.

Finally, to demonstrate the protective efficacy of the nasal vaccination, we challenged the i.n. or i.m. immunized hACE2 mice with the SARS-CoV-2 (2.5 x 10^4^ TCID_50_) (Extended Fig. 6a). Then, we measured viral titers in nasal turbinate, throat, lung, and brain tissues from the early to the middle phase of infection (1-4 dpi). Virus titers on 1-3 dpi in the nasal turbinates and throats of the animals received i.n. immunization were significantly lower than the virus titers in animals that received i.m. immunization (Extended Fig. 6b, c, *P* <0.05-0.0001), respectively. Contrary to the PBS group, hACE2 mice immunized with S-Fc via either i.n. or i.m. routes showed complete inhibition of virus growth and dissemination in the lung (Extended Fig. 6d) and brain (Extended Fig. 6e) at 2 and 4 dpi. Overall, these data suggest that the i.n. delivery of the S-Fc vaccine induces local humoral and cellular immune responses that provide more efficacious protection in the upper respiratory tract against SARS-CoV-2 infection than the i.m. route.

### Intranasal immunization with the S-Fc reduces transmission of SARS-CoV-2 between the immunized and unimmunized hamsters

SARS-CoV-2 is highly contagious and transmits among individuals mainly through the mouth or nose and possibly inhalation into the lungs of those who are near the infected. Hence, it is highly desirable that a vaccine can prevent or reduces SARS-CoV-2 transmission from vaccinated individuals to unvaccinated or vice versa. Hamster-to-hamster^23^ or hamster-to-human ^24^ transmission of SARS-CoV-2 occurs.

We first immunized hamsters and showed the hamsters that were i.n. immunized with S-Fc were protected from ancestral SARS-Cov-2 infection (Extended Fig. 7; Supplementary Information C). Hamsters that were either i.n. or i.m. immunized by the S-Fc (Extended Fig. 7a) developed significantly higher levels of IgG (Fig. 6a, p<0.001) and nAb (Fig. 6b, p<0.0001) in the sera in comparison with the PBS control hamsters. However, the hamsters that were immunized with the S-Fc via the i.n. route, compared to the i.m. route, had much higher levels of S-specific IgA Ab in both the nasal washes (Fig. 6c, p<0.01) and BAL (Fig. 6d, p<0.05). To test the transmissibility, we used a unidirectional airflow chamber. Naïve hamsters were exposed to the i.n. immunized hamsters that were infected with SARS-CoV-2, and vice versa (Fig. 6e). As a control, the naïve hamsters were also exposed to the infected naive hamsters. The i.m. immunized hamsters by the S-Fc were used as a control. Subsequently, we i.n. infected hamsters with a high titer of SARS-CoV-2 (1X10^5^ TCID_50_/hamster) to replicate a breakthrough infection (Fig. 6f). Exposure of naïve or immunized hamsters to airflow produced by infected immunized or naive hamsters was performed for 10-14 days.

**Figure 6.**
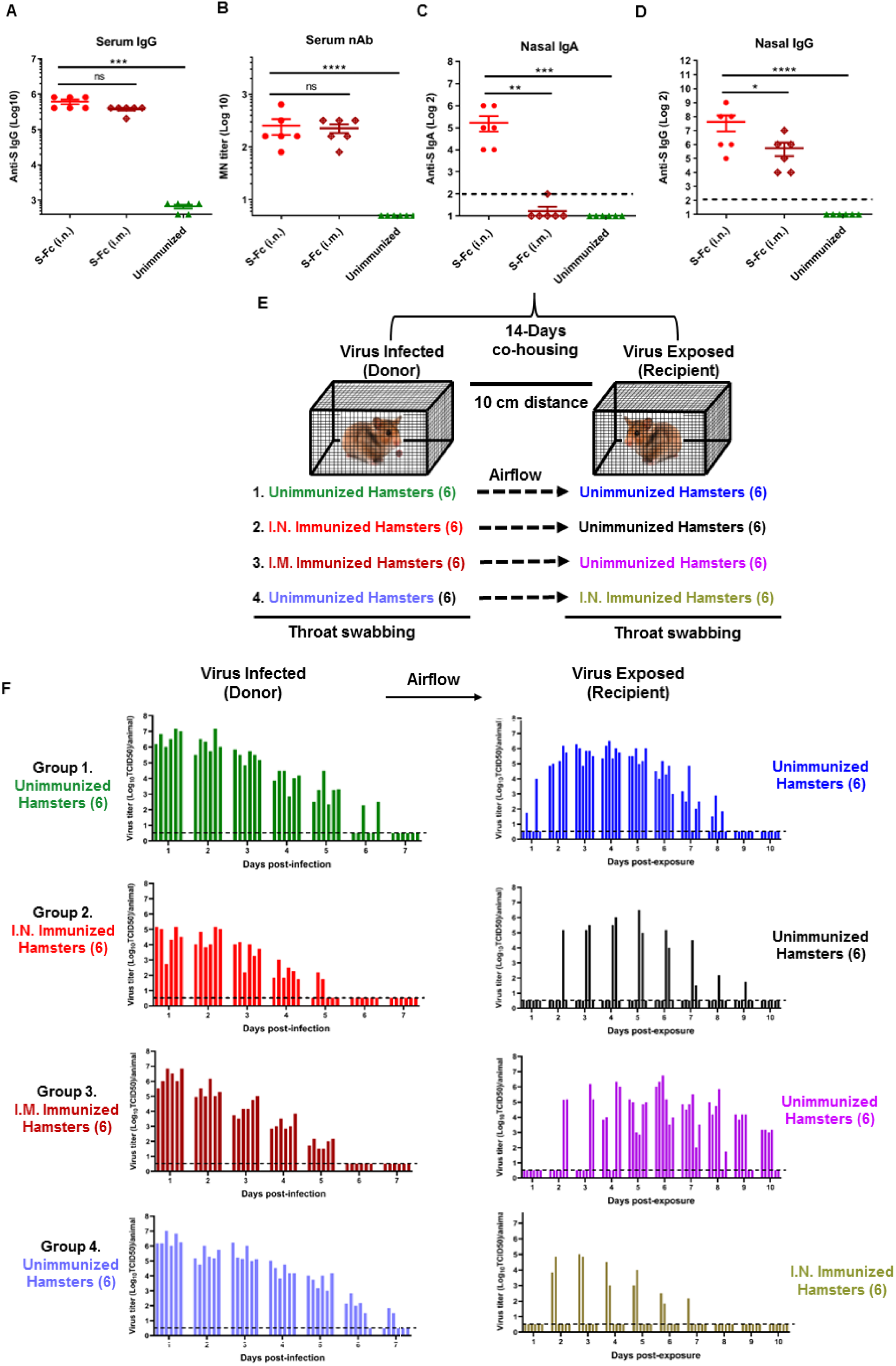
Intranasal immunizations with the S-Fc protein reduce virus transmission from vaccinated to unvaccinated hamsters or vice versa. **A+B+C+D.** Groups of male hamsters (n=6) were i.n. or i.m. immunized with 30 μg of S-Fc in combination with 30 μg of CpG twice in a 2-week interval. The unimmunized hamsters were used as a negative control. Anti-SARS-CoV-2 S-specific IgA, IgG Ab, and neutralizing Ab titers in sera and nasal washings were measured by ELISA 14 days after the boost. The data represent a geometric mean with 95% CI. One-way ANOVA with Dunnett’s multiple comparison tests was used for data analysis. **E.** A schematic illustration of the experimental design. Briefly, 4 groups of hamsters are included and each group has 6 hamsters for virus donors and 6 hamsters for virus recipients. The donor hamsters were either unimmunized or immunized with S-Fc plus CpG via i.n or i.m. routes. Then the donor hamsters were infected with 1 × 10^5^ TCID_50_ SARS-CoV-2. 14 hours later, donor hamsters in the wire cages were separately cohoused with 6 unimmunized hamsters or immunized hamsters within the same isolator. Throat swabbing was performed for 7 days for donor hamsters (1-7 dpi) and 10 days for recipient hamsters (1-10 days after exposure). **F**. Virus loads in the throat swabs from the donor and recipient hamsters. The presence of infectious live virus in the samples was measured in VAT cells for 4 days of culture. The viral titers were shown as TCID50 from each animal sample. Each bar represents one animal.

As expected, all six unimmunized hamsters exposed to the SARS-CoV-2-infected unimmunized hamsters had a high titer of live virus detected in their throats, which peaked at 3-6 dpi (Fig. 6f, group 1), indicating that SARS-CoV-2 infection and airborne transmissions were successful. All six i.n. immunized hamsters infected with the virus exhibited much less body-weight loss and had a lower level of live virus detection in their throat swabs (Fig. 6f, group 2) than the infected naïve hamsters (Fig. 6f, group1), verifying again that the i.n. immunizations with the S-Fc induced protective immunity at viral entry sites. In the exposed animals, four of six naïve unimmunized hamsters exposed to the i.n. immunized, SARS-CoV-2-infected hamsters had no detectable live virus during the 10 days (Fig. 6f, group 2). In contrast, all six naïve unimmunized hamsters exposed to the virally-infected hamsters that were i.m. immunized with the S-Fc protein displayed a high titer of live SARS-CoV-2 virus in their throats. However, the virus detection was delayed 2-3 days (Fig. 6f, group 3), suggesting i.m. immunized animals can spread the virus. Four of the six the i.n. immunized hamsters exposed to SARS-CoV-2-infected naïve unimmunized hamsters had no live virus detectable during the ten days (Fig. 6f, group 4). Although the live virus was detected in two of the i.n. immunized hamsters, the virus titers were significantly lower (Fig. 6f, group 4). Viral load was also quantified as SARS-CoV-2 N gene RNA in throat swab fluid on 1-4 dpi. The results from the N RNA detection were consistent with the results of the live-virus detection (Extended Fig. 8b-e). Together, our study unambiguously demonstrates that the i.n. immunization by the S-Fc provides protection by decreasing viral shedding and preventing airborne SARS-CoV-2 transmission (Extended Table 1) in a hamster model.

## Discussion

In this study, we explore a respiratory vaccination strategy by taking advantage of the mucosa-specific, FcRn-mediated IgG Ab transfer pathway, aiming at reducing or eliminating SARS-CoV-2 replication and spread in or through the nose and lung. The present study produced several lines of evidence. First, the S-Fc-immunized mice had higher IgG and IgA Abs in nasal washes, BAL, and blood, compared with those immunized by S alone or FcRn KO mice immunized by S-Fc. Also, the S-Fc-immunized mice exhibited stronger nAb activity relative to the control animals. Second, as opposed to the PBS control animals, hamsters or hACE2-transgenic mice i.n. immunized by S-Fc developed resistance to the ancestral SARS-CoV-2 infection, exhibiting reduced virus replication in the nasal turbinate, lung, and mouse brain. We detected high levels of live virus in the brain of the infected unimmunized hACE2 mice but not in the infected hamsters. This discrepancy may be explained by the expression level and localization of human ACE2 in transgenic animals. For example, hamsters expressing human ACE2 also have high virus titers detected in the brain following exposure to SARS-CoV-2^25^. Third, the S-Fc-immunized, hACE2 mice displayed excellent protection against infections by the SARS-CoV-2 Delta or Omicron variants. An additional booster is required to protect hACE2 mice from Omicron infection completely. However, this is not surprising because Omicron variants possess many mutations in its S protein and are shown to be resistant to immune responses elicited by the S protein from the ancestral strain. Likewise, to induce broad nAbs against the Omicron strain, it is evident that two boosters are also required for the mRNA vaccination ^26^. Fourth, remarkably, most S-Fc-immunized animals exhibited significantly reduced inflammation in the lungs of hACE2 mice. Finally, in the transmission study, in striking contrast to the PBS-immunized hamsters or hamsters immunized by the i.m. route, most i.n. S-Fc immunized hamsters that were infected with the virus did not spread the virus to unimmunized hamsters. Similarly, most i.n. s-Fc immunized hamsters were not infected with the virus spreading from the infected unimmunized hamsters. Our study demonstrates that respiratory mucosal immunization with S-Fc is essential to generating protective immunity against infection and transmission by the SARS-CoV-2 viruses in preclinical studies.

Several mechanisms may account for protecting against SARS-CoV-2 infection and transmission. FcRn bounds S-Fc in acidic pH conditions. A slightly acidic pH in the respiratory tract^27^ is expected to facilitate FcRn to retain antigens in airway mucosal surfaces and transfer them across the airway barrier, as shown by our previous studies^28, 29^ and others ^30^. The local Ab immune responses can represent a primary barrier of immune defense against viral infections of the respiratory tract, characterized by the presence of sIgA in the nasal mucosa or IgG in the BAL^10^. The natural SARS-CoV-2 infection induces mucosal sIgA, which dominates the early nAb response to SARS-CoV-2 with potent neutralizing activity^5, 31^. Furthermore, the levels of mucosal Abs determine the viral load and the amount of time required to recover from systemic symptoms. In this study, the mice and hamsters that were i.n. immunized by S-Fc developed high levels of IgA Abs in the nasal washes and BAL. These results are consistent with the findings of previous studies where FcRn-targeted respiratory immunizations against viral mucosal infections ^9, 28, 29^ and adenovirus-vectored SARS-CoV-2 nasal vaccine^32, 33^ elicit high Ab responses. In addition, the mice or hamsters i.m. immunized by S-Fc failed to produce significantly high levels of IgA Abs in the nasal washes, despite high IgG Abs in the BAL. A robust SARS-CoV-2 infection in nasal turbinates is readily detectable after the adoptive transfer of the nAbs^34^. This observation is further supported by our findings that hACE2 mice succumb to nasal infection following an adoptive transfer of serum Abs from i.n. immunized mice by S-Fc (Extended Fig. 9). Indeed, i.m. vaccination fails to significantly reduce viral load in nasal swabs or nasal turbinate ^35^. Considering SARS-CoV-2 also infects cells via a cell-to-cell fashion^36^, it may be difficult for serum IgG Abs to block viral entry and spread in the upper respiratory tract, or a high level of IgG Abs is required.

FcRn-mediated respiratory delivery of vaccine antigen facilitates the production of memory immune responses. The induction of S-specific memory immune responses is crucial for a vaccine to provide sustained protection^37^. The S- or RBD-specific nasal Abs can last for at least 9 months in COVID-19 patients^38^. FcRn-mediated respiratory vaccination with S-Fc induces and sustains high levels of S-specific IgA and IgG and plasma cells 8 months after the boost, which they resist lethal SARS-CoV-2 infection at least 6 months after the boost. TRM T cells in the lung can promote rapid viral clearance at the site of infection and mediate survival against lethal viral infections, especially in severe COVID patients^17, 39^. Our study demonstrated that a higher percentage of CD4^+^ or CD8^+^ TRM T cells were present in the lungs of mice that received i.n. S-Fc immunization than in those of PBS control animals. Furthermore, the i.n immunization induced significantly more lung-resident TRM cells than the i.m. route. This is consistent with previous findings by directly delivering viral antigens into the airway ^9, 17^. Additionally, FcRn prolongs IgG half-life^14, 15^. FcRn binding to S-Fc, thus, is expected to stabilize S-Fc in vivo, which would enhance FcγRs-mediated uptake of S-Fc by antigen-presenting cells^40, 41^. Likely, TRM T cells are particularly critical against the SARS-CoV-2 variants that undergo rapid mutations and evade antibody immunity. Further studies are needed to demonstrate the duration that these TRM T and B cells^16, 42^ may persist in the lung and how they contribute toward long-term protection against the SARS-CoV-2 variants. SARS-CoV-2-specific CD8^+^ TRM cells persist for at least two months after viral clearance in the nasal mucosa^39^.

Several rationales could explain the rise in breakthrough infections, two of which are most likely. First, i.m. vaccination couldn’t effectively induce local immunity in the respiratory mucosa. Indeed, the i.n., but not i.m., immunizations with S-Fc induce nasal IgA and TRM T cells in the lung. Second, individuals infected by SARS-CoV-2 through the nasal epithelial cells can asymptomatically shed the infectious virus for an unknown time. Hence, we envision the nasal vaccination as an essential complementary strategy to the i.m. vaccines that are authorized or undergoing clinical trials. Because FcRn can transfer the immune complexes across the mucosal epithelial barrier with or without pre-existing Abs^13, 43^, we do not expect that the efficacy of the FcRn-targeted mucosal vaccination is influenced by pre-existing SARS-CoV-2 immunity. The FcRn-targeted nasal vaccine can be used for an unimmunized first dose or booster for those who have been immunized, even previously infected by SARS-CoV-2. The combination of mucosal and parenteral vaccines has been proven effective at mucosal entry against SARS-CoV-2 infections^7, 9, 44, 45^.

In addition to pneumonia and acute respiratory distress, some symptomatic or asymptomatic COVID-19 patients, including those with breakthrough infections, are reported to experience a long COVID ^46, 47^. The exact causes of the long COVID-19 are elusive; they may involve direct effects of viral infection or indirect influences on the brain. In humans, the nasal olfactory epithelium expresses a high level of ACE2 receptor. Although it is debatable, SARS-CoV-2 may enter the brain by crossing the neural-mucosal interface in olfactory mucosa ^47^ or infecting the olfactory epithelium or bulb, or both^48^. Indeed, SARS-CoV-2 is identified to replicate in brain astrocytes^49^. Alternatively, inflammatory cytokines derived from inflamed lungs or other organs may cross the blood-brain barrier^50^. Consequently, a virus or cytokine may cause encephalitis or necrosis, leading to a long COVID. The hACE2 mice are highly susceptible to SARS-CoV-2 infection with the virus detected in the brain^20^. Our study confirms this fact and further shows that FcRn-mediated respiratory vaccination can prevent brain viral infection in hACE2 mice. Likely, the sIgA or TRM T cells induced by S-Fc can efficiently block the viral replication in nasal turbinates, resulting in reduced viral infection in the olfactory epithelium or access to the olfactory bulb.

Taken together, we developed a respiratory vaccine using pre-stabilized SARS-CoV-2 S antigen that stimulates local and systemic protective immunity. Our working model states that FcγR-expressing mucosal antigen-presenting cells, including dendritic cells, take up the FcRn-transported spike and subsequently migrate to mediastinal lymph nodes where they prime the CD4^+^ T cells and initiate the cognate B cell response in the germinal centers. By increasing the persistence of the S-Fc in tissue and circulation, the vaccination may further enhance the development of long-term humoral and cellular immunity. The substantially longer half-life of IgG in humans compared to mice or hamsters (21 vs. 6 days) would ensure high and persisting levels of S-Fc in human immunization. Also, the S-Fc and human IgG1 had a lower affinity to the mouse FcγRI, but it exhibits a high affinity with human FcγRI (Fig. 1d), suggesting our S-Fc vaccine may work more efficiently in humans. Although the i.n. administered viral vaccines induce protection in the respiratory tracts^32, 33^, a protein-based nasal vaccine may be preferred, especially in young, elderly, and immunocompromised populations. Together, our results indicate that FcRn-mediated respiratory immunization can be an effective and safe strategy for maximizing the efficacy of vaccinations against infection and transmission of SARS-CoV-2 and its emerging variants.

## Materials and Methods

### Cells, Antibodies, and Viruses

Vero E6 (with high expression of endogenous ACE2, Cat No. NR-53726) and VAT (Vero E6-TMPRSS2-T2A-ACE2, Cat No.NR-54970) were from Biodefense and Emerging Infections Research Resources Repository (BEI Resources, Manassas, VA). Chinese hamster ovary (CHO) cells were purchased from the American Tissue Culture Collection (ATCC, Manassas, VA). Vero E6, VAT, and CHO cells were maintained in complete Dulbecco’s Minimal Essential Medium (DMEM) (Invitrogen Life Technologies), both supplemented with 10% fetal bovine serum (FBS), 2 mM L-glutamine, nonessential amino acids, and antibiotic and antifungal (100 units/ml of penicillin, 100 μg/ml of streptomycin, and 250 ng/ml of amphotericin B). Vero E6, VAT, and CHO cells routinely tested negative for *Mycoplasma* sp. by real-time PCR. Recombinant CHO cells were grown in a complete medium with G418 (Invitrogen, 1 mg/ml). All cells were grown at 37°C in 5% CO_2_.

S-specific mAbs 2TP1B11, 2TP2C7, and 2TP22E7 were procured from the BEI. S specific mAbs 40150-R007, 40952-MM57, and SPD-M128, as well as anti-S polyclonal antibodies 40590-T62 and 40150-T62-Cov2, were acquired from Sino Biologicals. Hamster IgG2 was purchased from BD (Cat # 553294). Motavizumab, an antibody against the respiratory syncytial virus (RSV) F protein, was acquired from Cambridge Biologics (Brookline, MA). The sera from the convalescent COVID-19 patients or health persons were a gift from the Biotech Laboratories (Rockville, MD). The horseradish peroxidase (HRP)-conjugated anti-human IgG (Cat# 2081-05), anti-goat IgG (Cat# 6160-05), streptavidin (Cat# 7100-05), and biotin-labeled goat anti-mouse IgA (Cat# 1040-08) were obtained from Southern Biotech (Birmingham, AL). HRP-conjugated anti-mouse IgG Fab (Cat# A9917) and anti-human IgG Fab (Cat# SAB4200791) were from Sigma. HRP-conjugated anti-mouse IgG (Cat# PA1-28568) and anti-hamster IgA (sab3003a) were obtained from Invitrogen (Waltham, MA) and Brookwood Biomedical (Jemison, AL), respectively. Goat anti-hamster IgG (Cat# NB1207141) was acquired from Novus (Centennial, CO). We purchased recombinant biotinylated human and biotinylated mouse FcRn/β2m (Cat# FCM-H82W4 and FCM-M82W6), and biotinylated human and mouse FcγRI proteins (Cat# FCA-H82E8 and CD4-M82E7), and biotinylated human ACE2 protein (Cat# AC2-H82E6) from AcroBiosystems (Newark, DE). Human C1q protein was a gift from Dr. Sean Riley (Complement Technology, Cat # A099). Mouse C1q protein (Cat# M099) was procured from Complement Technology (Tyler, TX).

SARS-CoV-2 ancestral strain hCoV-19/USA/NY-PV08410/2020 (abbreviated as NY strain, Cat# NR-53514), Delta strain hCoV-19/USA/PHC658/2021(B.1.167.2) (Cat# NR-55611), and Omicron strain USA/MD-HP20874/2021(B.1.1.529) (Cat# NR-56461) were obtained from BEI Resources with the permission of the Centers for Disease Control and Prevention (CDC). Viruses from the BEI were passed in either Vero E6 (for ancestral and Delta strains) or VAT cells (for the Omicron strain). At 72 hr post-infection, tissue culture supernatants were collected and clarified before being aliquoted and stored at −80 °C. The virus stock was respectively titrated by using the Median Tissue Culture Infectious Dose (TCID_50_) assay. All virus experiments were performed in an approved Animal Biosafety Level 3+ (ABSL-3+) facility at the University of Maryland using appropriate powered air-purifying respirators (PAPR) and protective equipment.

### Mice and Golden Syrian hamsters

The Institutional Animal Care and Use Committee approved the animal protocol at the University of Maryland (#R-APR-20-18 and #R-MAR-21-19). The animals were acclimatized at the animal facility for 4–6 days before initiating experiments. Animals from different litters were randomly assigned to experimental groups, and investigators were not blinded to allocation during experiments and outcome assessment. Wild-type (WT) C57BL/6 mice were purchased from Charles River Laboratories (Frederick, MD), and FcRn KO mice were a gift from Dr. Derry Roopenian (Jackson Laboratory, Bar Harbor, ME). Transgenic mice expressing human ACE2 by the human cytokeratin 18 promoter (K18-hACE2) represent a susceptible rodent model^20, 51^.

Specific-pathogen-free, 6–8-weeks-old, female and male B6.Cg-Tg(K18-ACE2)2Prlmn/J (Stock No: 034860, K18-hACE2) hemizygous C57BL/6 mice and control C57BL6 mice (non-carriers) were purchased from the Jackson Laboratory and used for breeding pairs to generate pups for research. All the offspring were subjected to genotyping, and only the hemizygous K18-hACE2 mice were chosen for future use. Seven-week-old male/female *Golden Syrian* hamsters were obtained from Charles River Laboratories. Animals have been maintained in individually ventilated cages at ABSL-2 for noninfectious studies or in isolators within the ABSL-3 facility for studies involving SARS-CoV-2 viruses. Immunizations and virus inoculations were performed under anesthesia. We avoided using volatile chemical anesthetics known to increase the permeability of the respiratory epithelial barrier in nasal immunizations. All mice were anesthetized with an intraperitoneal (i.p.) injection of fresh Avertin at 10-12.5 μl of working solution (40 mg/ml) per gram of body weight -and laid down in a dorsal recumbent position to allow for recovery. Hamsters were sedated with Dexmedetomidine (50-250 μg/kg) via subcutaneous injection for immunization and virus infection, or they were anesthetized with isoflurane for collecting blood, nasal washes, and throat swabs.

### Construction of plasmids expressing a trimeric and prefusion-stabilized SARS-CoV-2 spike (S) and Spike-Fc (S-Fc)

The entire amino acid (aa) sequence corresponding to the S protein of the ancestral SARS-CoV-2 strain Wuhan-Hu-1 was retrieved from Genbank (*MN908947.3*). We designed an S gene of SARS-CoV-2 with some modifications described below and synthesized it from GenScript (*Piscataway, NJ*). The S protein precursor has two well-defined cleavage sites: S1/S2 and S2’. To produce a non-cleavable S protein, we performed mutagenesis at the cleavage site S1/S2 (R685A) and S2’(R816A) of the S gene to keep the S protein in a pre-cleavage conformation. On the surface of coronaviruses, S glycoprotein exists predominantly in the prefusion form^52^. A prefusion structure of SARS-CoV-2 S is critical for maximizing immunogenicity. To produce a prefusion form of S antigen, we made a pair of point mutations to proline (K986P and V987P) as previously described^52^, which prevents the formation of a helix associated with the post-fusion conformation (Fig. 1A). The S protein also naturally exists as a trimer on the virions or virally infected cells^53^. To facilitate the trimerization of soluble S protein, we engineered a foldon domain from T4 bacteriophage fibritin protein^18^ to the C-terminus of the extracellular domain of the S (residues 1-1213) gene to facilitate the trimerization. To target FcRn for delivery, we selected human IgG1 Fc. The rationale for using human IgG1 is consistent with the fact that it has the highest affinity for activating FcγRI, but the lowest affinity for inhibiting FcγRIIB^54^. In IgG1 Fc, the complement C1q-binding motif was eliminated (E318A/K320A/K322A)^55^ (Fig. 1A) to prevent the binding of human and mouse C1q. Because human IgG1 Fc typically forms a disulfide-bonded dimer, we created a monomeric Fc by substituting cysteines 226 and 229 with serine (C226S and C229S) to eliminate the disulfide bonds, as we reported previously ^9^. The modified S gene (named by S-Fc) encodes the extracellular domain of the S protein, followed by a trimeric motif (foldon) and a monomeric human IgG1 Fc segment. The S-Fc gene was further codon-optimized for optimal expression in CHO cells and synthesized and cloned into eukaryotic expression plasmid pcDNA3.1 via *Kpn* I and *Xho* I sites to generate the recombinant plasmid pcDNA3.1-S-Fc (Fig. 1A).

A control plasmid, pcDNA3.1-S, was produced by replacing the human IgG1 Fc portion with the 6x His tag sequence. For this purpose, an inverse PCR was performed using pcDNA3.1-S-Fc as a template, and the primer pairs: 5’-CACCTTCCTGGGCCATCATCACCATCACCATTGACTCGAGTCTAGAGGGCCCG-3’ and 5’-ATGGTGATGGTGATGATGGCCCAGGAAGGTGGACAGCAGCACCCACTCGCCAT-3’. After the transformation of the inverse PCR segment into the competent *E. coli* strain, homologous recombination allows for the generation of the circular plasmid pcDNA3.1-S, which encodes a trimeric soluble S protein with the foldon domain and 6x His tag.

### Generation of plasmids expressing hamster FcRn/β2m or FcγRI

To construct plasmid for the expression of hamster FcRn and β2m, the DNA sequences encoding the extracellular domain of hamster FcRn (1-300 aa) or full-length hamster β2m were firstly amplified from hamster lung tissue by RT-PCR. The parimer paris for hamster FcRn were 5’-GCGGGTACCGCCACCATGGGGATGCCCCAGCCC-3’, 5’-TATCTCGAGTTACTCGTGCCACTCGATCTTCTGGGCCTCGAAGATGTCGTTCAGGCC GTGGTGATGGTGGTGATGGTGATGCGAAGATCTGGCTGGAGCA-3’. The primer pairs for hamster β2m amplification were 5’-TATGTCGACGCCACCATGGCTCGCTCCGTGGCCG-3’,5’-GCGTCTA GACTATTTTTCGAACTGCGGGTGGCTCCACATGTCTCGTTCCCAGGTGAC-3’. The corresponding FcRn cDNA sequence comprises a coding sequence for 8x His tag and Avi tag at the 3’ end for protein purification and site-specific biotinylation, respectively, while the β2m cDNA included a Strep II tag at its 3’ end. Then, the FcRn and β2m cDNAs were subsequently cloned into a dual-expression vector pBud-CE4.1 (Invitrogen) via the *Kpn* I/*Xho* I sites (for FcRn) and *Sal* I/*Xba* I sites (β2m) to generate a plasmid pBud-haFcRn/β2m.

To create a plasmid expressing hamster FcγRI, an RT-PCR was performed to amplify the gene sequence encoding the ectodomain of FcγRI from hamster lung tissue. The primer pairs were 5’-GCGGGTACCGCCACCATGTGGCTCCTAACAACCCT G-3’ and 5’-TATCTCGAGTTACTCGTGCCACTCGATCTTCTGGGCCTCGAAG ATGTCGTTCAGGCCGTGGTGATGGTGGTGATGGTGATGAGGGCCTGATGACTGAGG AC-3’. The tandem 8x His tag- and Avi tag-encoding sequence at the 3’ end of the FcγRI segment was designed for the purpose of affinity purification and site-specific biotin labeling, as described above. Next, the hamster FcγRI segment was digested with *Kpn* I and *Xho* I and inserted into the pBud-CE4.1 vector to generate a plasmid pBud-haFcγRI.

### Expression and characterization of the prefusion-stabilized S and S-Fc fusion proteins

The pcDNA3.1 plasmids encoding S and S-Fc were transfected into CHO cells using PEI MAX 40000 (Fisher Scientific, Cat# NC1038561). Stable cell lines were selected and maintained under G418 (1 mg/ml). Expression and secretion of the S or S-Fc fusion proteins were determined by immunofluorescence assay, SDS-PAGE, and Western blotting analysis. We produced the soluble S or S-Fc proteins by culturing CHO cells in a complete medium containing 5% FBS with ultra-low IgG. The proteins were captured by Protein A column (ThermoFisher Scientific, Cat# 20356) for the S-Fc protein or Histidine-tagged Protein Purification Resin (R&D Systems, Cat # IP999) for the S protein, eluted with 0.1M Glycine (pH 2.5), and neutralized with 1M Tris-HCl (pH8.0). Glycine and Tris-HCl in the protein solution were replaced with PBS three times using centrifugation with Amicon Ultra-15 Centrifugal Filter Unit (50K) (Millipore, Cat# UFC905024). Protein concentrations were determined using a NanoDrop spectrophotometer (Thermo Scientific).

### Expression and purification of the hamster FcRn/β2m and FcγRI proteins

To express hamster FcRn/β2m or FcγRI proteins, 293T cells were transfected with recombinant plasmid pBud-haFcRn/β2m or pBud-haFcγRI. The supernatants from the cell culture were harvested after 48 hrs and loaded onto an anti-His resin column and eluted with 0.1M glycines (pH 2.5). The eluted proteins were neutralized with 1M Tris-HCl (pH 8.0) and the protein buffer was replaced with PBS by using Amicon Ultra-15 Centrifugal Filter Unit (10K). The purified proteins were visualized by SDS-PAGE and Coomassie staining. The site-specific biotinylation was then conducted by using BirA biotin-protein ligase standard reaction kit (Cat# BirA500) from Avidity Biosciences (Aurora, CO).

### SDS-PAGE gel and Western blotting

Protein quality was assessed by 8-15% SDS-PAGE gel under reducing conditions. Proteins in gels were either stained with Coomassie blue dye in gel or used for transfer onto nitrocellulose membranes (GE Healthcare). The membranes were blocked with 5% milk in PBST (PBS and 0.05% Tween-20) and incubated with appropriate primary and HRP-conjugated secondary antibodies, as indicated in the Figure legends. The immobilon Western chemiluminescent HRP substrate (Millipore, Cat# WBKLS0100) was used to visualize protein bands in membranes and images captured by the Chemi Doc XRS system (BioRad).

### Samplings of nasal washes, bronchoalveolar lavage (BAL), and throat swabs

The nasal washes and BAL fluids were collected from the mouse under euthanasia with over-dosage of Avertin. For sampling nasal washes, we removed the lower jaw. A small incision was made over the ventral aspect of the trachea; then, a syringe with a blunt plastic tip was inserted into the trachea toward the nasal cavity. A total of 1 ml PBS was gently injected into the nasopharynx and collected when it flowed from the external nares. For BAL collection, the syringe was inserted into the trachea but toward the lungs, and 1 ml of PBS was carefully injected into the lungs by keeping the syringe in position. The PBS was retrieved back to obtain BAL fluids. The nasal washes and BAL fluids were centrifuged to remove cellular debris, concentrated to 350 μl with Amicon 0.5 ml centrifugal filter unit (10K) (Millipore, Cat# UFC501096), and the supernatants were stored at -20°C.

To collect nasal washes in hamsters, animals were anesthetized with isoflurane and small-sized feeding needles (20G) were used to inject 500 l sterile PBS into the nostrils (250 μl each side). The outflows were collected in a petri dish as expelled by the hamster. The volume was increased to 0.5 ml with the addition of cold PBS. To collect throat swabs from the anesthetized hamsters, the swab was first moistened in 650 μl DMEM media with 1% inactivated FBS and then placed into the throat, where it was gently rubbed around ten times. These swabs were then soaked for 5 minutes in the vials containing the remainder of the media. Subsequently, the throat swabs were removed, and the samples were vortexed and stored at −80°C for further virological analysis.

### Enzyme-linked immunosorbent assay (ELISA) and Enzyme-linked immunosorbent spot (ELISpot)

For the detection of SARS-CoV-2 S-specific antibodies IgA and IgG in nasal washes, BAL fluid, and sera collected along the detection timeline after the immunizations, 96-well plates (Maxisorp, Nunc) were coated with 1 µg/ml of the S protein as described above in 100 l coating buffer (PBS, pH7.4) per well and incubated overnight at 4°C. Plates were then washed four times with 0.05% Tween 20 in PBS (PBST) and blocked with blocking buffer (2% bovine serum albumin in PBST) for 2 hr at room temperature. The serially diluted specimens (nasal wash, BAL, or sera) from animals were added to each well and incubated for 2 hrs. After washing six times with PBST, the detection antibodies were added and incubated for 1.5 hr at room temperature. HRP-conjugated rabbit anti-mouse IgG (1: 20,000, Invitrogen, Cat# PA1-28568) was used for measuring mouse IgG, while biotin-labeled goat anti-mouse IgA Ab (1:5000, Southern Biotech, Cat#1040-08) plus HRP-conjugated streptavidin (1:7000, Southern Biotech, Cat# 7100-05) were used for measuring mouse IgA antibody. To detect S-specific hamster IgG, goat anti-hamster IgG (1:4000, Novus, Cat# NB1207141) plus HRP-conjugated rabbit anti-goat IgG (1:5000, Southern Biotech, Cat# 61-6065) were used. To determine S-specific hamster IgA, HRP-conjugated anti-hamster IgA (1:250, Brookwood Biomedical, Cat# sab3003a) was used. One hundred microliter TMB (tetramethylbenzidine) (BD, Cat# 555214) was used as a substrate to visualize the signals. Reactions were stopped with 100 μl of 1 M sulfuric acid. Optical density at 450 nm was determined using a Victor III microplate reader (Perkin Elmer). Titers represent the reciprocal of the highest dilution of samples showing a 2-fold increase over the average OD_450_ nm values of the blank wells.

The ELISA assays were also used to measure interactions of the S-Fc or S with 1) human, mouse, or hamster FcRn/β2m heterodimer (ACROBiosystems, Cat# FCM-H82W4 for human FcRn/β2m; Cat # FCM-M82W6 for mouse FcRn/β2m; at the acidic pH (6.0) and neutral pH (7.4) conditions; 2) human, mouse, or hamster FcγRI (ACROBiosystems, Cat# FCA-H82E8 for human FcγRI; Cat# CD4-M82E7 for mouse FcγRI); 3) human ACE2 (ACROBiosystems, Cat# AC2-H82E6); and 4) human and mouse C1q protein (Complement Technology, Cat#A099 and #M099). To facilitate detection, all FcRn/β2M heterodimer, FcγRI, ACE2, and C1q proteins were conjugated with biotin. In brief, ELISA plates were coated with S-Fc or S protein in PBS (1 μg/well for FcRn/β2M binding or 200ng/well for FcγRI and hACE2 binding) overnight at 4°C. After blocking for 2 hr, the 2-fold serial diluted target proteins (4-4000 ng/ml of FcRn/β2m, 0.4-400 ng/ml of FcγRI and hACE2) were added and incubated for 2 hr at room temperature. For the C1q binding assay, S-Fc or S proteins were used to coat plates at a serial dilution (800-7.8 ng/well), and a biotin-conjugated human or mouse C1q (2 μg/ml) was used for detection. For all assays, the streptavidin-HRP was from Southern Biotech (1:5000) and TMB was used to visualize the colorimetric signals.

For measuring S-specific antibody-secretin cells (ASCs) in bone marrow, an ELISpot kit (MabTech, Cat# 3825-2H) was used. The 96-well ELISpot plates (MabTech, Cat# 3654-TP-10) were pre-wetted with 35% ethanol and washed five times with sterile water plus 1 time with PBS. The plates were then coated with S protein at 20 µg/ml overnight at 4°C (100 l/well) and blocked with RPMI 1640 complete medium with 10% FBS for 2 hr at room temperature. Bone marrow cells from femurs and tibias were collected in RPMI 1640, filtered through a 70 m strainer, and subjected to ACK lysis. Serial dilutions of single-cell suspensions were prepared in RPMI 1640 and added to the coated wells for 18-24 hr at 37°C in 5% CO_2_. After incubation, the plates were emptied and washed five times with PBS, then incubated with biotinylated anti-mouse IgG (0.5 μg/ml) for 2 hr at room temperature. After washing with PBS, HRP-streptavidin (1:700) was added and incubated for 1 hr. The samples were developed with TMB substrate until distinct spots emerged. After washing, the plates were stored upside down in the dark to dry overnight at room temperature. Spots were counted with an ELISpot reader (AID, Germany).

### Quantification of SARS-CoV-2 virus and viral RNA

The number of infectious virus particles in the specimen of ancestral SARS-CoV-2 or Delta strain infected animals was determined in Vero E6 cells by 50% tissue culture infectious dose (TCID_50_) endpoint dilution assay as described^56^. The quantification of the Omicron strain was performed in VAT cells. To increase the sensitivity, VAT cells were also used in detecting ancestral viruses and Omicron variants in the throat swab samples, as the overexpression of the hACE2 and TMPRSS2 in VAT cells enhances the replication efficiency of the SARS-CoV-2^57^. Briefly, cells were plated at 15,000 cells/well in DMEM with 10% FBS and incubated overnight at 37°C with 5.0% CO_2_. Media was aspirated and replaced with DMEM with 1% inactivated FBS for virus infection. Animal tissues including nasal turbinate, lung, brain, intestine, and kidney were homogenized in the TissueLyser LT (Qiagen). After centrifuging at high speed (14000 rpm, 10 min), the 10-fold serial dilutions of supernatants were used to infect the cell monolayers in 96 well plates, and the CPE was checked after four days. Positive (virus stock of known infectious titer) and negative (medium only) controls were included in each assay. The virus titer was expressed as TCID_50_/ml (50% infectious dose (ID_50_) per milliliter) by using the Reed-Muench method^56^.

To monitor viral RNA levels in virus-infected animal samples (throat swabs), total RNAs were isolated by using the PureLink RNA mini kit (Invitrogen, Cat# 12183018A) and subjected to the one-step quantitative real-time reverse transcription-PCR assay (qRT-PCR) using TaqMan Fast Virus 1-Step Master Mix (ThermoFisher, Cat# 4444432) as described previously^58, 59^. A SARS-CoV-2 nucleocapsid (N) specific primers and probe sets were used: Forward primer: 5’-GACCCCAAAATCAGCGAAAT-3’; Reverse primer: 5’-TCTGGTTACTGCCAGTTGAATCTG-3’; and probe: 5’-FAM-ACCCCGCATTACGTTTGGTGGACC-BHQ1-3’). Briefly, viral RNA was expressed as N gene RNA copy numbers from each swab or animal, based on an RNA standard included in the assay, which was created via the *in vitro* T7-DNA-dependent RNA transcription of a linearized DNA molecule containing the target region of the N gene full-length by using MEGAscript T7 Transcription Kit (ThermoFisher, Cat# AM1334), and purified with MEGAclear Transcription Clean-Up Kit (ThermoFisher, Cat# AM1908). The amplifications of qRT-PCR were performed with a CFX96 Touch Real-Time PCR System (Bio-Rad) using the following conditions: reverse transcription at 50 °C for 5 minutes, initial denaturation at 95 °C for 20 s, then 40 cycles of denaturing and annealing/extending at 95°C for 3 s and 60 °C for 30 s. The lower limit of detection (LOD) was 10^1.5^ copies per reaction.

### Microneutralization (MN) assay

Neutralizing antibodies were measured by a standard microneutralization (MN) assay on Vero-E6 (for ancestral and Delta strains) or VTA cells (for Omicron strain) as previously described^15^. The sera were heat-inactivated at 56°C for 30 min and followed by 2-fold serial dilution, after which the diluted sera were incubated with 100 TCID_50_ of SARS-CoV-2 virus (ancestral, Delta, and Omicron strains) for 1 hr at 37°C, respectively. The virus-serum mixtures were added to Vero-E6 or VAT cell monolayers in 96-well plates and incubated for 1 hr at 37°C. After removing the mixture, DMEM with 1% inactivated FBS was added to each well and incubated for four days at 37°C for daily CPE observation. Neutralizing Ab titers are expressed as the reciprocal of the highest serum dilution preventing the appearance of CPE.

### Immunizations of mice and Golden Syrian hamsters and SARS-CoV-2 challenge

Six to eight-week-old female/male C57BL/6 mice, FcRn KO mice, and K18-hACE2 transgenic mice were intranasally (i.n.) immunized with 10 μg S-Fc, equal molar of recombinant S, or PBS in 10 μg CpG adjuvant (ODN1826, Invivogen, Cat# vac-1826-1) in a total volume of 20 μl. Previous studies showed that CpG does not increase the permeability of the airway respiratory barrier; in contrast, it does enhance the tight junction integrity of the bronchial epithelial cell barrier^60^. For intramuscular (i.m.) immunizations, mice were injected bilaterally in the quadriceps femoris with a 50 μl volume containing 10 μg S-Fc antigen in 10 μg CpG. Six to eight-week-old female/male hamsters (n = 6-8 per group) were vaccinated via i.n. or i.m. route with an 80 μl volume containing 30 μg S-Fc and 30 μg CpG. The mice or hamsters were boosted with the same vaccine formulations two or three weeks later.

Two or three weeks after the boost, blood was collected from each animal; 3 days later, the animals were transferred to the ABSL-3+ facility for virus challenge. The K18-hACE2 mice were i.n. infected with lethal doses of SARS-CoV-2 virus in a total volume of 50 µl (2.5 x 10^4^ TCID_50_ ancestral SARS-CoV-2 and Delta strains, or 1 x 10^6^ TCID_50_ for Omicron strain). Due to their high susceptibility, the aged K18-hACE2 mice were challenged with 5 x 10^3^ TCID_50_ ancestral strain. The hamsters were i.n. infected with 1 x 10^5^ TCID_50_ of ancestral SARS-CoV-2 strain with a final volume of 100 µl. After infection, animals were monitored daily for morbidity (weight loss), mortality (survival), and other clinical signs of illness for 14 days.

Animals losing above 25% of their body weight following infection or reaching the humane endpoint were humanely euthanized.

At the indicated time points after the virus infection, nasal washes or throat swabs were sampled to monitor the virus shedding from the upper respiratory tract. To further measure the virus replication and tissue lesion *in vivo*, 50% of the animals in each group were euthanized at 4 or 5 dpi and different organs and tissues, including nasal turbinate, trachea, lung, brain, heart, and intestine, were harvested. The left lung lobe was fixed in a 10% neutral buffered formalin solution for histopathology analyses, while the right lung lobes and other tissues were homogenized in DMEM by Tissue Lyser (Qiagen). The homogenates were cleaned by centrifugation (15000 rpm for 10 minutes), and supernatants were collected to measure viral load.

### Lung pathology

To examine the lung pathology, lungs were removed from mice in each group and fixed in 10% neutral buffered formalin solution three days before transferring the tissues out of the ABSL-3 facility. The lungs were then paraffin-embedded, sectioned in five-micron thickness, and stained with Hematoxylin and Eosin (H & E) by Histoserv Inc (Germantown, MD). Stained lung sections were scanned using a high-definition whole-slide imaging system (Histoserv, Germantown, MD).

To determine the level of pulmonary inflammation, the lung inflammation was evaluated and scored by a board-certified veterinary pathologist blinded to the experimental design. A semi-quantitative scoring system, ranging from 0 to 5, was used to assess the following parameters: alveolitis, parenchymal pneumonia, inflammatory cell infiltration, peribronchiolitis, perivasculitis, and lung edema^61^. The inflammatory scores are as follows: 0, normal; 1, very mild; 2, mild; 3, moderate; 4, marked; and 5, severe. An increment of 0.5 was assigned if the inflammatory score fell between two.

### Intravascular labeling and flow cytometry

To discriminate the tissue-resident memory T cells (TRM) in the lung from the circulating T cells in the blood, the S-Fc immunized, or PBS control mice were anesthetized and intravenously injected with 3 μg of PE-CD3 Ab in 100 μl PBS through the retro-orbital route. After 5 min labeling, the treated mice were euthanized and bled. As described previously, single cells were isolated from the lung^28^. The lungs were perfused with 10 ml PBS, minced with scissors, and incubated in a digestion solution (RPMI with 1mg/ml of collagenase IV, 5 mM of CaCl2, 10 μg/ml DNase I) for 45-60 min at 37°C on a rotating rocker. The digestion was stopped with 5 mM EDTA, and then the cells were filtered through a 70 μm cell strainer and treated with ACK buffer to lyse the red blood cells. Next, cells were purified via the 30% Percoll centrifugation to get rid of most epithelial cells and cell debris. After washing and resuspension with PBS, cells were stained with a Fixable Live/Dead Yellow staining dye (Invitrogen, Cat# 501121527) to differentiate between live and dead cells. Single-cell suspensions were incubated with Fc block (anti-mouse CD16/CD32, 1 μg for 1x10^6^ cells, BD Biosciences, Cat# 553142) at 4°C for 30 min. After washing with FACS buffer (2% FBS and 2mM EDTA in PBS), cells were stained with FITC-CD3 Ab (BD Bioscience, Cat# BDB555275), APC-H7-CD4 Ab (BD Bioscience, Cat#590181), BV421-CD8 Ab (Biolegend, Cat#100753), APC-CD69 Ab (BD Bioscience, Cat#BDB560689), and PE-eFluor610-CD103 Ab (eBioscience, Cat# 61103182) for 1 hr at 4°C. After washing, cells were fixed with 2% paraformaldehyde for 20 min and resuspended in 350 ul 1% BSA/PBS for phenotyping by FACSDivacytometer (BD Biosciences). To set up the compensation control, Abc total antibody compensation beads (Invitrogen, Cat# A10497) and Arc amine reactive compensation beads (Invitrogen, Cat# A10346) were used. The acquired data were analyzed using the FlowJo software (Tree Star).

### Airborne transmission experiments in hamsters

Airborne transmission of the SAS-CoV-2 is more efficient than fomite transmission in hamsters^62^. Hence, we examined the capacity of the i.n. immunization to reduce airborne transmission of the ancestral SAS-CoV-2 between animals. The i.m. immunized hamsters were used as controls. To this end, the hamsters were either i.n. or i.m. immunized with S-Fc or left unimmunized as described above. All hamsters were single-housed in stainless steel wire cages in an isolator, where they were grouped into donors (infected) and recipients (sentinel). The isolators provided a unidirectional airflow from the donors to the recipients at an air speed of 78 L/min. The airborne transmission study was conducted following the booster. All donor hamsters at anesthesia were i.n. inoculated with 1 × 10^5^ TCID_50_ of SARS-CoV-2 (100 μl) and placed upstream of the airflow location in an isolator. 12 hours later, the recipient hamsters were placed downstream of the airflow in the same isolator. The donor and recipient cages were seated at a distance of 10 cm to avoid direct contact with animals and the effect of dust particles generated from the bedding material. After infection, the body weight and clinical signs of the hamsters were monitored daily for 14 consecutive days. Throat swabs were sampled daily for seven days in donor hamsters, while the recipient hamsters were sampled for ten days as the onset of infection in the recipients was a few days later than that in the donors.

### Statistics analysis

All data were analyzed with the Prism 9.0 software (GraphPad). The student *t*-test was used to compare the means between two groups, while One-way ANOVA was used to compare the difference if three or more groups were involved. Meanwhile, a Post Hoc test was applied after One-way ANOVA. Dunnett’s multiple comparisons test was used to compare means from different treatment groups against a single control group. The Turkey test was performed to identify the difference between the two groups. To compare the Kaplan-Meier survival curves, the Mantel-Cox test was used. Fisher’s exact test was conducted for comparisons of transmission capacities among various groups. All statistical methods used in each experiment are indicated in the Figure legends. The level of statistical significance was assigned when P values were < 0.05. The statistical significance was further classified as four levels: * (P<0.05), ** (P<0.01), *** (P<0.001), and **** (P<0.0001).

## Acknowledgments

We are grateful to BEI Resources for supplying us with the SARS-CoV-2 virus and its variants, cells, and antibodies. We thank Dr. Chinta Lamichhane who shared the sera from the convalescent COVID-19 patients and healthy persons. We are most grateful for the technical help from Ms. Ely Elena Guevara and Mr. Takele Yazew. This work was supported in part by NIH grants AI146063, AI130712 (X.Z.), and MAES grants from the University of Maryland (X.Z.). W.T. and A.R. are supported by the intramural research program of the USDA ARS.

## Author Contributions

W.L., and X.Z. designed experiments and analyzed data. W.L., W. T., and X.Z. wrote the paper. W. L., T.W., A.R., Z. M., H.Y., G.A., and T.L. conducted experiments. W.T. provided critical materials, interpreted data, and made editorial suggestions.

