## Supplemental data for "An FcRn-targeted mucosal vaccine against SARS-CoV-2 infection and transmission"

### Supplementary Information (SI)

#### A. Intranasal vaccination with the S-Fc protein protects against SARS-CoV-2 in the aged K18-hACE2 transgenic mice.

The mortality and fatality of COVID-19 are highly skewed toward older adults and age is negatively correlated with immune responses after vaccination. The aged mice develop more severe lung damage than the young adult mice upon SARS-CoV-2 infection or re-infection (*63*). To assess the immune response and protective efficacy of our vaccine in aged mice, we performed i.n. immunization with two doses of S-Fc (10 μg/mouse) at a 2-week interval (Extended Fig. 4a) and used PBS-immunized mice as controls. As shown in Extended Fig. 4b, although high SARS-CoV-2-specific IgG was elicited in all S-Fc immunized mice two weeks after the boost, the levels of serum IgG antibodies produced in the aged mice were generally lower than those of the young adult mice. After the immunized mice received a second boost (*Fig. 4a*), the levels of serum IgG and neutralizing Ab elicited in the aged mice were comparable to those of the young adult mice that received 2-times i.n. immunizations (Extended Fig. 4b, c). We i.n. challenged the immunized aged mice with ancestral SARS-CoV-2 virus ($5 \times 10^3$ $TCID_{50}$). All aged mice in the PBS control group displayed high susceptibility to virus infection and suffered fast weight loss, resulting in 100% death. In contrast, the S-Fc-immunized aged mice did not exhibit obvious body-weight loss and clinical signs (Extended Fig. 4d) and all mice were fully protected, leading to 100% survival after the challenge (Extended Fig. 4e). Strikingly, no live virus can be measured in the throat swabs of the S-Fc-immunized mice throughout the infection period of 1-6 dpi in comparison to the active virus replication in the throat samples from the PBS control mice. (Extended Fig 4f *1-4 dpi: P<0.0001; 5 dpi: P<0.001;*

*6 dpi: P<0.01*). These results indicated the complete blockage of virus amplification and shedding from the airways of the infected animal by S-Fc immunization after three times immunization.

**B. Intranasal vaccination with the S-Fc protein elicits durable protection in K18-hACE2 transgenic mice**

Although the authorized booster vaccinations in adults elicit high levels of neutralizing Abs against the SARS-CoV-2, antibody levels can wane substantially 3-4 months after vaccination [64]. To address whether the i.n. immunization with the S-Fc can sustain long-term protection (Extended Fig. 4g), we measured the serum IgG and neutralizing Abs in the hACE2 mouse sera six months after the boost. The majority of the immunized mice maintained a significant level of S-specific IgG Ab (Extended Fig. 4h, $p<0.001$) and the neutralizing Ab activity (Extended Fig. 4i, $p<0.01$) in their sera compared to PBS control animals, although the levels were lower than those in the immunized sera collected two weeks following the boost. To investigate long-term protection, we further challenged mice with ancestral SARS-CoV-2 virus ($2.5 \times 10^4$ $TCID_{50}$) six months after the boost (Extended Fig. 4j). Following the challenge, mice immunized with the S-Fc exhibited significantly reduced viral replications in nasal turbinate, lung, and brain tissues (Extended Fig. 4j), while the PBS control mice displayed a considerably higher level of viral replications. Overall, intranasal delivery of the SARS-CoV-2 vaccine engendered an effective and long-term memory immune response and provided sustained protection against challenges.

**C. Intranasal vaccination with the S-Fc protein results in protection against SARS-CoV-2 in Golden Syrian hamsters**

Golden Syrian hamsters are highly susceptible to the SARS-CoV-2 virus and exhibit disease symptoms similar to humans, including severe lung inflammation[61]. We showed that hamster FcRn bound human IgG1 and the S-Fc (Fig. 1C). Hence, hamsters were i.n. immunized with 30 µg of the S-Fc protein or PBS together with 30 µg CpG and boosted 2 weeks later (Extended Fig. 7a). As shown in Extended Fig. 7, the S-Fc immunized hamsters induced significantly higher levels of IgG (Extended Fig. 7b, $p<0.0001$) and neutralizing Ab (Extended Fig. 7c, $p<0.0001$) titers in comparison with the PBS-immunized hamsters. Next, we investigated whether hamsters i.n. immunized by the S-Fc resist to SARS-CoV-2 infection. Groups of hamsters (8 hamsters/group) administered intranasally with S-Fc or PBS were challenged with 1x $10^6$ $TCID_{50}$ of ancestral SARS-CoV-2 virus 17 days after the boost. Hamsters within the PBS group underwent mild to moderate body weight loss (Extended Fig. 7d), but none of them succumbed to viral infection. In contrast, the S-Fc-immunized hamsters did not show weight loss (Extended Fig. 7d). Meanwhile, the S-Fc immunized hamsters had significantly reduced the number of viruses in nasal washes compared to that of PBS-immunized mice at 2 and 4 dpi (Extended Fig. 7e, $p<0.01$-$0.0001$), suggesting the decline of virus shedding from these animals. Third, we measured viral replications in the nasal turbinate, trachea, lung, brain, and intestine 5 dpi (Extended Fig. 7f). High titers of live SARS-CoV-2 virus were detected in the nasal turbinate and lung tissues of the PBS-immunized control hamsters; on the contrary, no live virus was isolated from the nasal turbinate and lung tissues of animals who received i.n. immunization with the S-Fc. Hence, virus titers in the nasal turbinates and lungs of the animals that received the i.n. immunization on day 5 post-infection were significantly lower than the virus titers in the animals that received PBS at the corresponding time postinfection (Extended Fig. 7f, $P < 0.0001$) for the virus titers in the nasal turbinates and lungs, respectively. In addition, we did not find a live virus in the trachea and brain tissues of either PBS- or the S-Fc-immunized hamsters. These data clearly indicate that the i.n. immunization with S-Fc provides protective immunity against SARS-CoV-2 infection in hamsters.

### Extended Data Figure/Table Legends

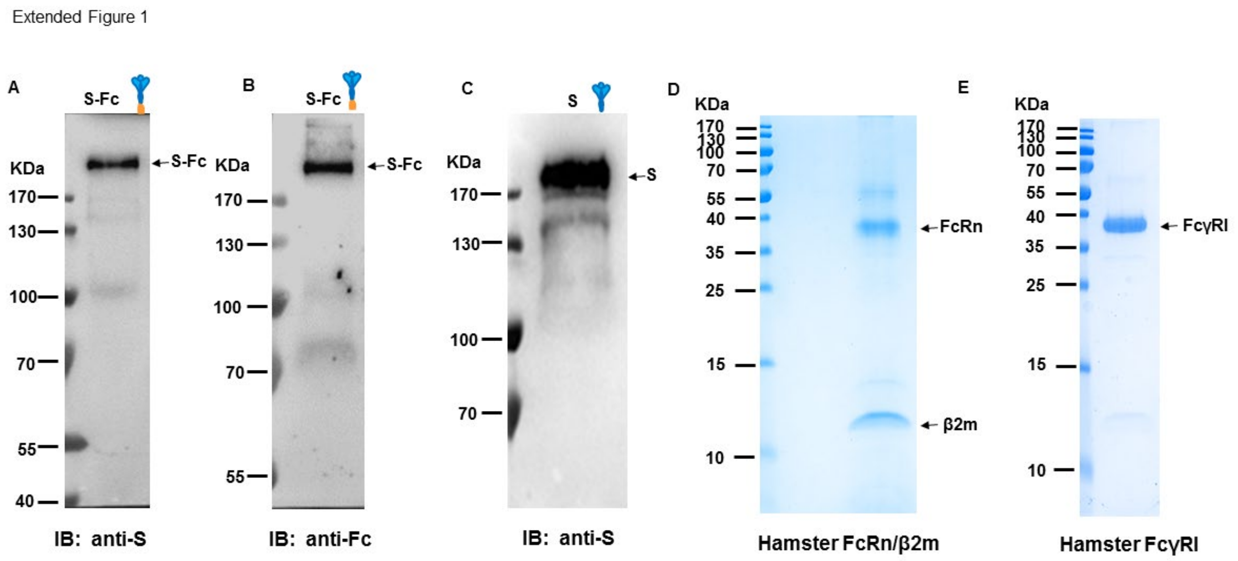


**Extended Data Fig. 1. Expressions of S-Fc, S, hamster FcRn/b2m, and FcγRI proteins.** The S-Fc fusion protein (**A+B**) or S (**C**) protein purified from the stable CHO cell line was identified by Western blot. The S-Fc or S proteins were subjected to SDS-PAGE and Western blot analyses and detected by either anti-S (**A+C**) or goat anti-human IgG-Fc primary Abs (**B**). The S-Fc or S proteins were visualized by HRP-conjugated secondary Abs and the ECL method. The purified hamster FcRn/β2m proteins (**D**) or hamster FcγRI proteins (**E**) from the recombinant plasmid-transfected 293T cells weres visualized by coomassie blue staining.

Extended Figure 2

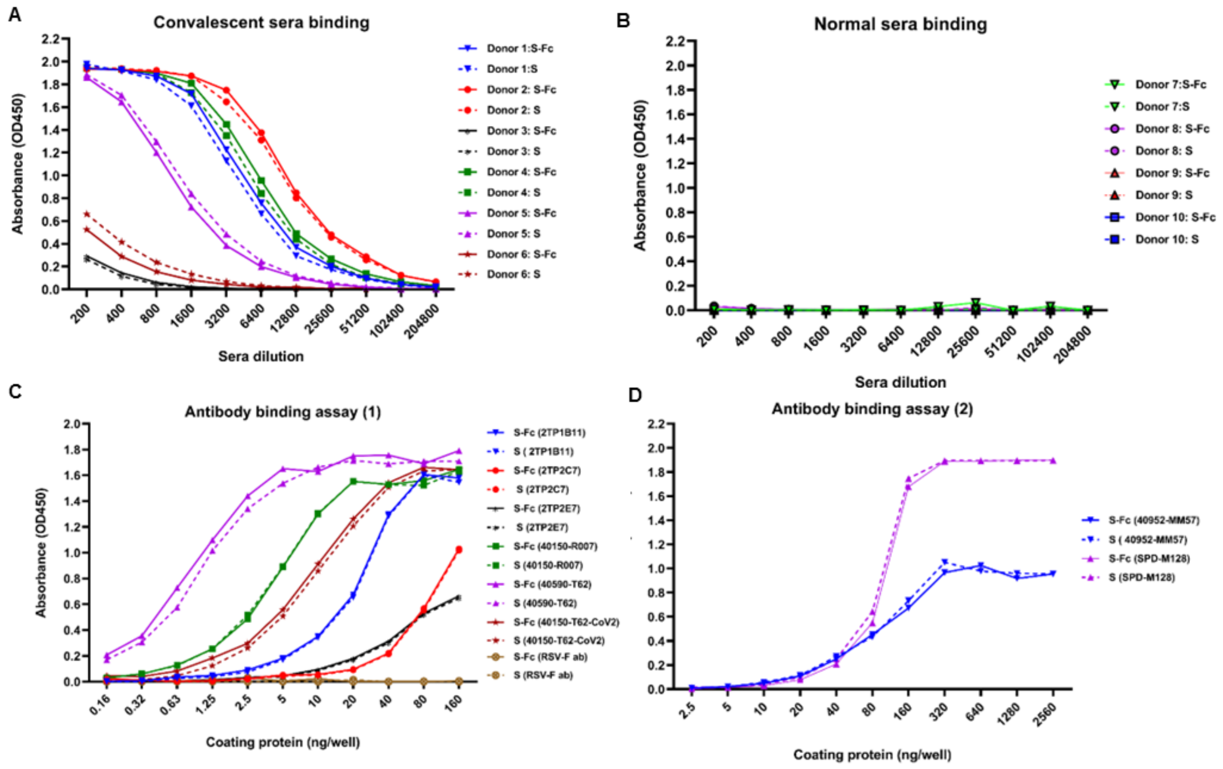


**Extended Data Fig. 2. Interactions of the purified S or S-Fc with SARS-CoV-2 S-specific antibodies.** Interactions of the purified S or S-Fc with COVID-19 convalescent human serum (**A**), normal human serum (**B**), and a set of SARS-CoV-2 S-specific mAbs (**C+D**). The specific binding was detected by the ELISA method.

Extended Figure 3

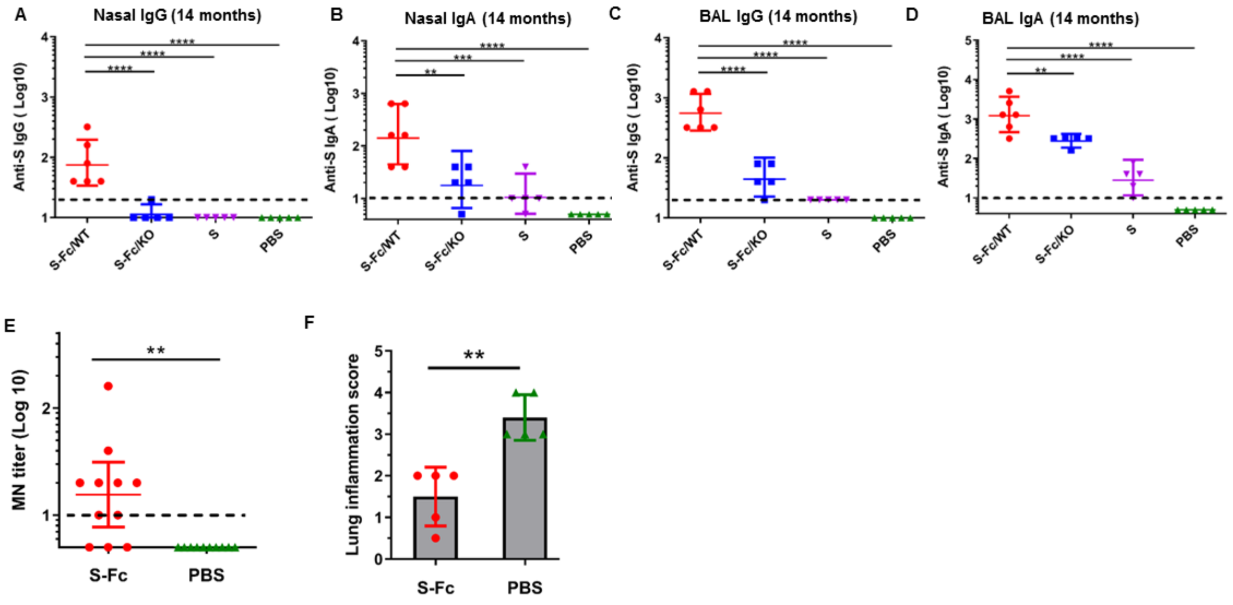


**Extended Data Fig. 3. Anti-S-specific Ab titers and lung inflammation score**. Anti-S-specific Ab titers in nasal washings (**A+B**), and BAL (**C+D**) after 14 months of the boost. SARS-CoV-2 S-specific Abs were measured by ELISA from 5-6 representative mouse samples in each group. The data represent a geometric mean with 95% CI. One-way ANOVA with Dunnett's multiple comparison tests was used for data analysis. (**E)** SARS-CoV-2 Delta-specific nAbs in human ACE2 mice after the boost. The nAb activity in the sera was determined by the micro-neutralization test. The data represent a geometric mean with 95% CI. (**F)** After the Delta strain challenge, the inflammatory responses of each lung section were scored *in a blind manner.* Statistical differences were determined by the student *t*-test.

Extended Figure 4

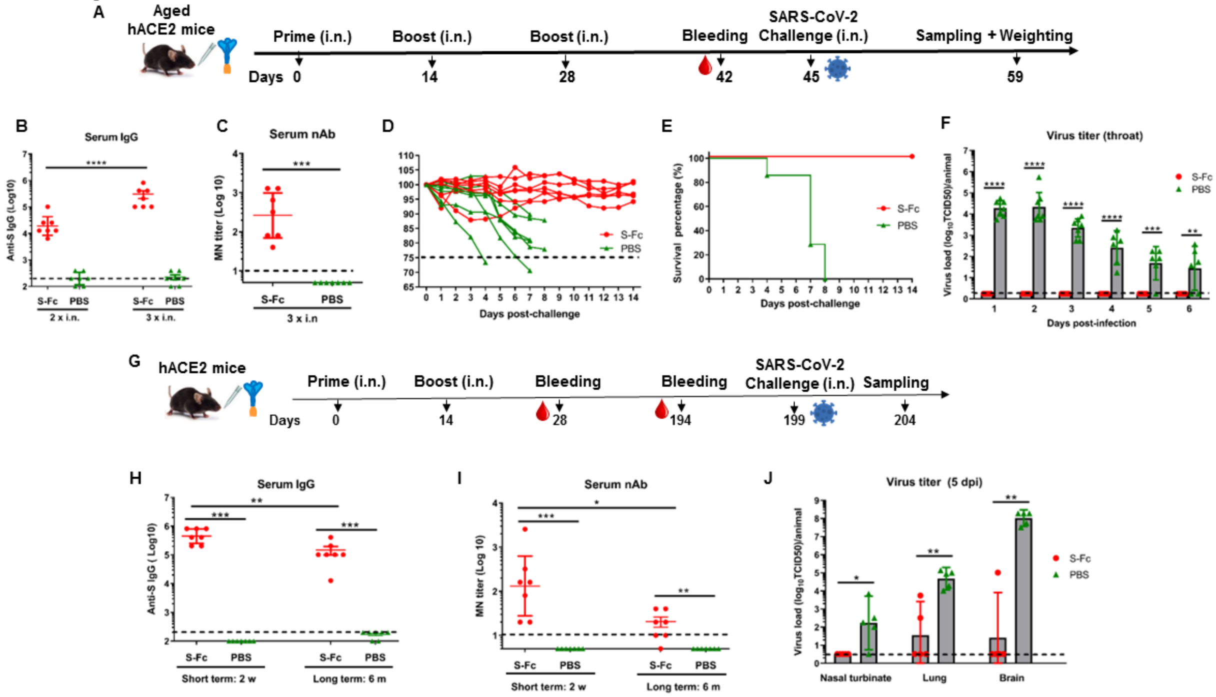


**Extended Data Fig. 4. Intranasal immunizations with the S-Fc induces protection in aged mice and protective memory immune response.**

**A.** Ten µg of S-Fc, or PBS in combination with 10 µg of CpG was i.n. administered into the age-matched 12-18 months-old human ACE2 mice. Mice were boosted twice at a 2-week interval following primary immunization. Bleeding, virus challenge, and sampling were performed at the indicated time points.

**B**. Anti-S-specific IgG Ab titers in the aged mouse sera. The S-specific Ab titers were measured by coating with S protein in ELISA 14 days after the first and second time boosts. The IgG titers were measured in 7 representative mouse sera. 2 x: one-time boost; 3 x: two-times boost.

**C.** The nAb titer in the aged mouse sera. Two weeks after the second boost, sera sampled from 7 mice per group were heat-inactivated and serially diluted two-fold in PBS. The ancestral SARS-CoV-2 (100 $TCID_{50}$) was added and incubated at 37°C for 1 hr. The mixture was added to Vero-E6 cells and incubated at 37°C for 96 hr, the nAb titers were determined and expressed as the reciprocal of the highest dilution preventing the appearance of the CPE.

**D**. Body-weight changes following the challenge. 17 days after the second boost, the aged mice (S-Fc group, n=7; PBS group, n=7) were i.n. challenged with ancestral SARS-CoV-2 virus (5 X $10^3$ $TCID_{50}$) and weighed daily for 14 days. Mice were euthanized at the end of the experiment or when a humane endpoint was reached.

**E.** Survival following SARS-CoV-2 virus challenge in the aged mice. The percentage of aged mice protected on the indicated days is calculated as the number of mice that survived divided by the total number of mice in each group, as shown by the Kaplan-Meier survival curve.

**F.** Throat samples were collected daily for 7 days in each aged mouse after the challenge. The presence of infectious live virus was measured in VAT cells after 4 days of culture. The viral titers were shown as $TCID_{50}$ from each animal swab.

**G.** Ten µg of S-Fc or PBS in combination with 10 µg of CpG was i.n. administered into 8-week-old hACE2 mice. Mice were boosted once in a 2-week interval following primary immunization. Six months after the boost, mice were i.n. challenged with the ancestral SARS-CoV-2 virus. Blood collection, virus challenge, and sampling were performed at the indicated time points.

**H**. Anti-SARS-CoV-2 S-specific IgG Ab titers in sera two weeks (2 w) or six months (6 m) following the boost. The S-specific Ab titers were measured by coating the plates with S protein in ELISA. The IgG titers were measured in 7 representative mouse sera for each group. The data represent a geometric mean with 95% CI.

**I.** SARS-CoV-2 nAb in sera from the immunized mice (n=7) was determined using the $TCID_{50}$ test 2 weeks or 6 months after the boost. The nAb titers are expressed as the reciprocal of the highest dilution preventing the appearance of the CPE.

**J**. Viral titers in the nasal turbinate, lung, and brain 5 days after ancestral SARS-CoV-2 (2.5 X $10^4$ $TCID_{50}$) challenge. Supernatants of the nasal turbinate, lung, and brain homogenates were added to Vero-E6 cells and incubated for 4 days. The viral titers were shown as $TCID_{50}$ from each swab. The student *t*-test was used for the statistical assay.

Extended Figure 5

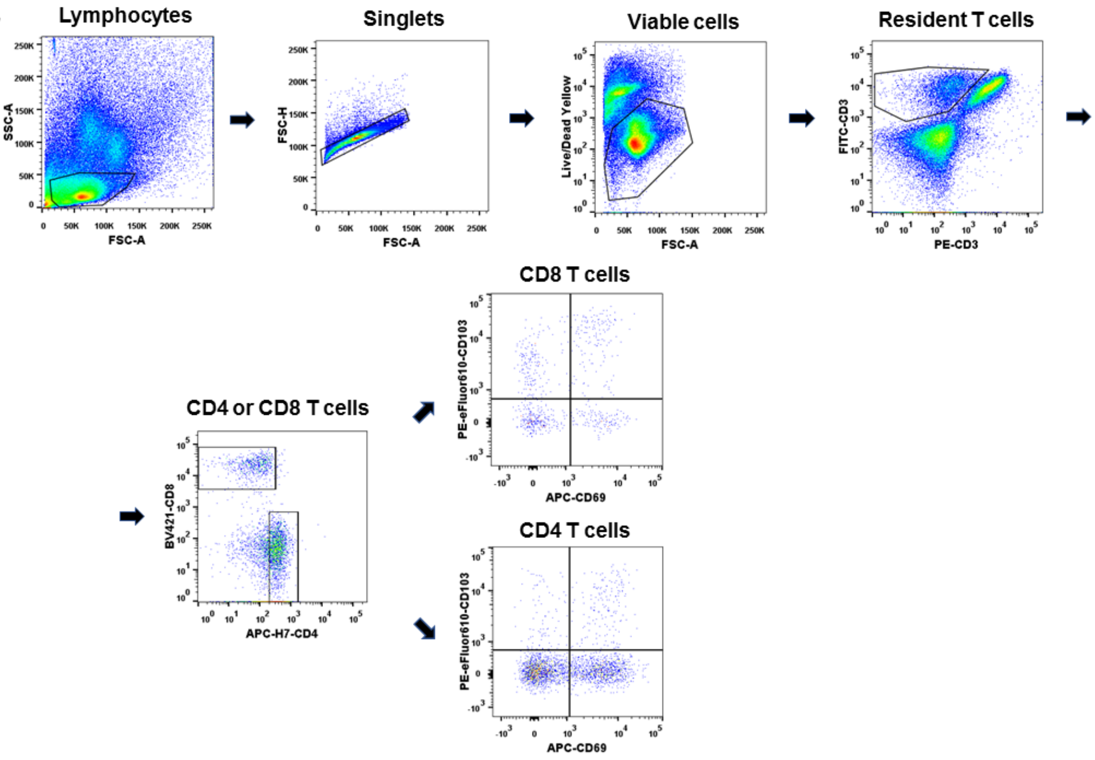


**Extended Data Fig. 5. Gating strategy for the identification of TRM T cells in the lungs**. The lymphocyte population was first gated from the lung tissue's mixed cells in the plot of FSC vs SSC. Then an FSC-H vs. FSC-A plot was used to select singlets, followed by a viability dye staining to exclude the dead cells. To distinguish resident cells from others, an intravenous (IV) staining strategy using PE-CD3 was applied to rule out the positively stained, circulating T cells. The negative population (IV-) showing the positive reaction for the in-vitro FITC-CD3 staining was characterized as a T subset located in the lung. This subset was further separated into CD4+ and CD8+T cells, and the representative phenotypes of TRM were defined as the co-expression of CD69 and CD103.

Extended Figure 6

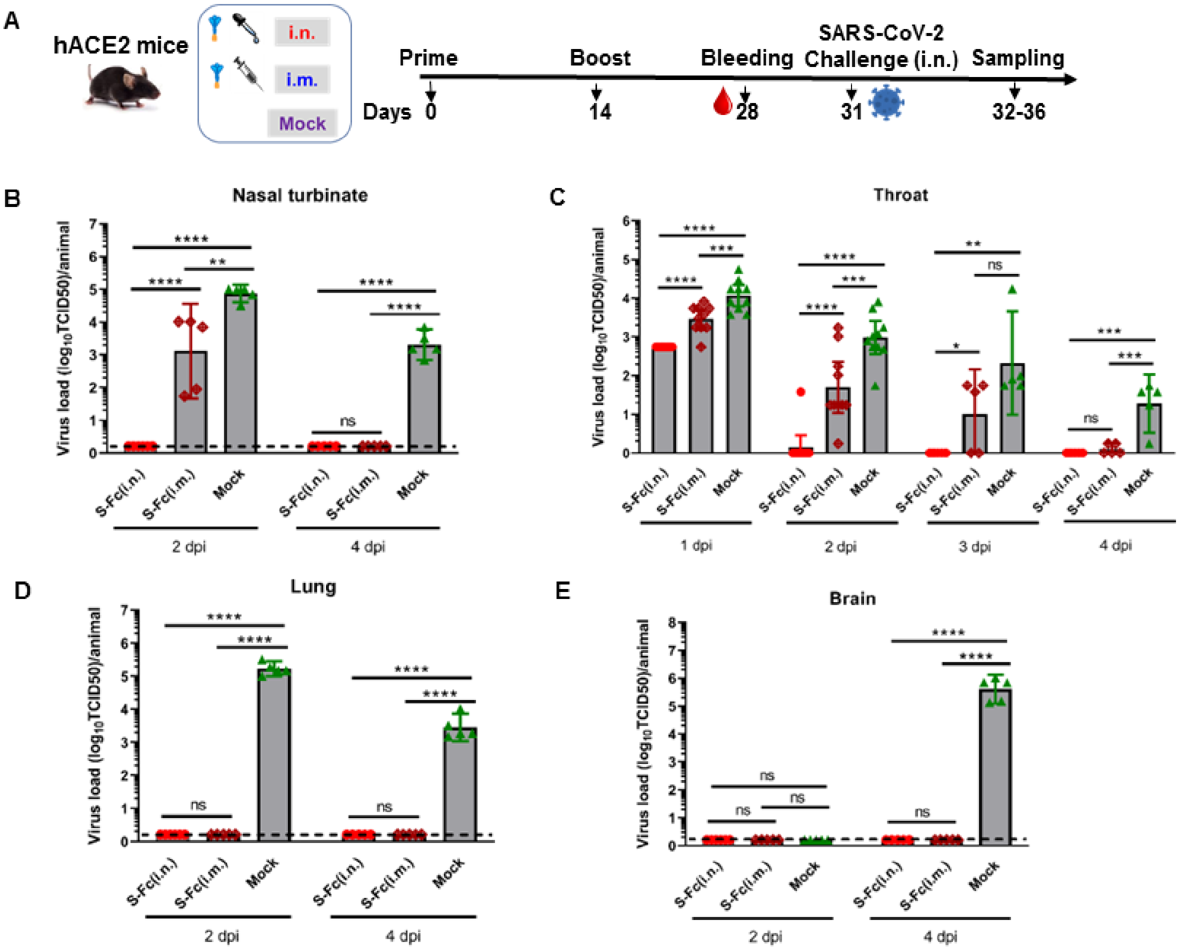


**Extended Data Fig. 6. FcRn-mediated intranasal vaccination significantly reduces viral replications in the upper respiratory tract.**

**A.** Ten µg of S-Fc with 10 µg of CpG was i.n. or i.m. administered into 6-8 week-old hACE2 mice (n=10-11). Mice were boosted 14 days after primary immunization. A group of mice (n=10) that were mock immunized as a negative control. Throat swabbing was performed daily from 1-4 dpi. Half mice in each group were euthanized at 2 and 4 dpi, respectively, for harvesting tissues and titrating the virus.

**B+C+D+E**. Samples were collected from throat swabs, nasal turbinates, lung, and brain tissues at multiple time points after the challenge, as displayed at the bottom. The live virus titers were determined by calculating $TCID_{50}$ values. One-way ANOVA followed by Turkey multiple comparisons test was used for the statistical assay.

Extended Figure 7

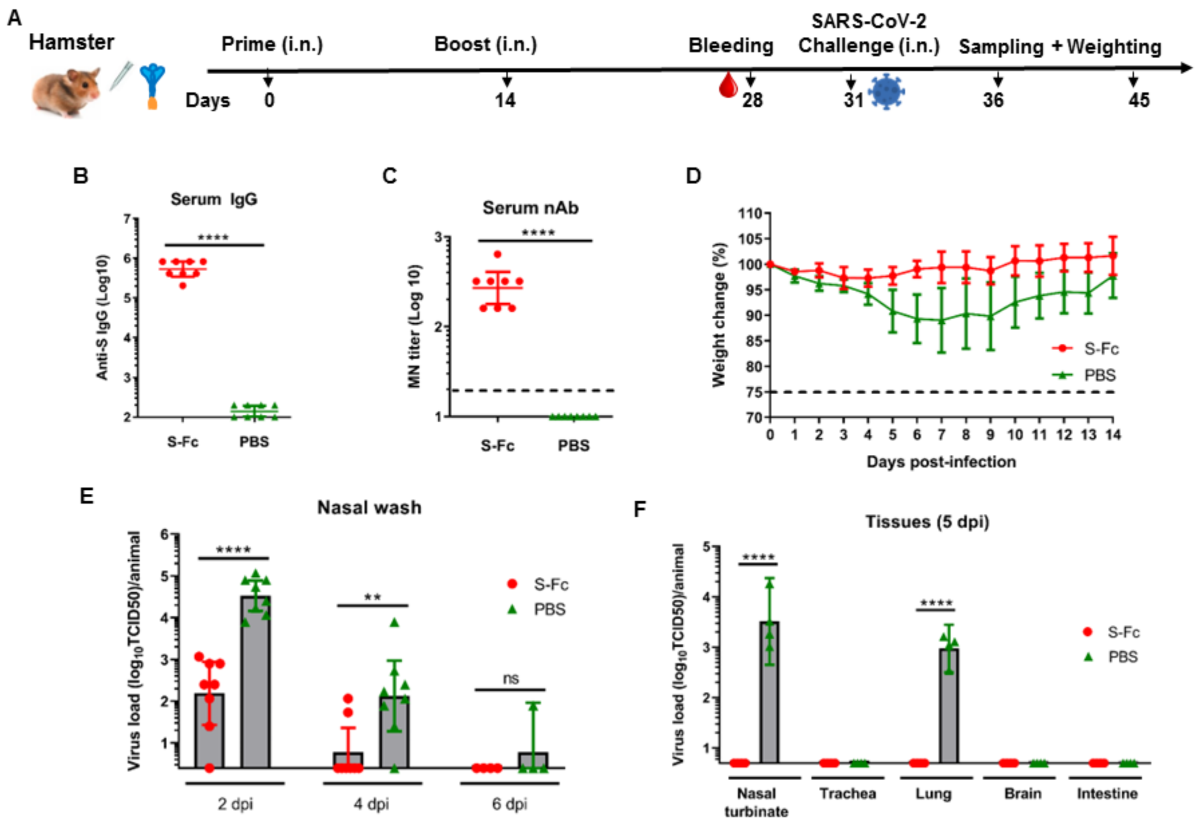


**Extended Data Fig. 7**. **Intranasal immunizations with the S-Fc protein protect *Golden Syrian* hamsters from SARS-CoV-2 infection.**

**A.** 30 μg of the S-Fc (n=8), or PBS (n=8) in combination with 30 μg of CpG was i.n. administered into female hamsters twice in a 2-week interval. Animals were challenged with ancestral SARS-CoV-2 ($1 \times 10^5$ $TCID_{50}$) and nasal washings were collected at 2, 4, and 6 dpi. Four hamsters from each group were euthanized and sampled at 5 dpi for titrating the virus. The 4 hamsters in each group were used for weight loss and survival assay.

**B.** Anti-SARS-CoV-2 S-specific IgG Ab titers in the hamster sera. The S-specific Ab titers were measured by coating with S protein in ELISA 14 days after the boost. The IgG titers from eight hamster sera per group were measured.

**C.** The nAb in the immunized hamster sera. Two weeks after the boost, sera sampled from eight hamsters per group were heat-inactivated and serially diluted two-fold in PBS. The neutralizing Ab activity in the sera was determined by the micro-neutralization test.

**D**. Changes in body weight after virus challenge. Seventeen days after the boost, hamsters were i.n. challenged with $1 \times 10^5$ $TCID_{50}$ of ancestral SARS-CoV-2 and weighed daily for 14 days. Hamsters were euthanized at the end of the experiment or when a humane endpoint was met.

**E.** Shedding of SARS-CoV-2 virus in nasal wash samples of the immunized and controlled hamsters. The virus levels were determined by a $TCID_{50}$ assay.

**F.** The viral titers in the nasal turbinate, trachea, lung, brain, and intestine at 5 dpi. Four animals from each group were euthanized for virus titration 5 days after the challenge. Supernatants of the nasal turbinate, trachea, lung, brain and intestine homogenates were added onto Vero-E6 cells and incubated for four days. The viral titers were shown as TCID50 from each animal swab. The student t-test was used for statistical analysis. A nonparametric test was applied if the data were not from the normally distributed population.

Extended Figure 8

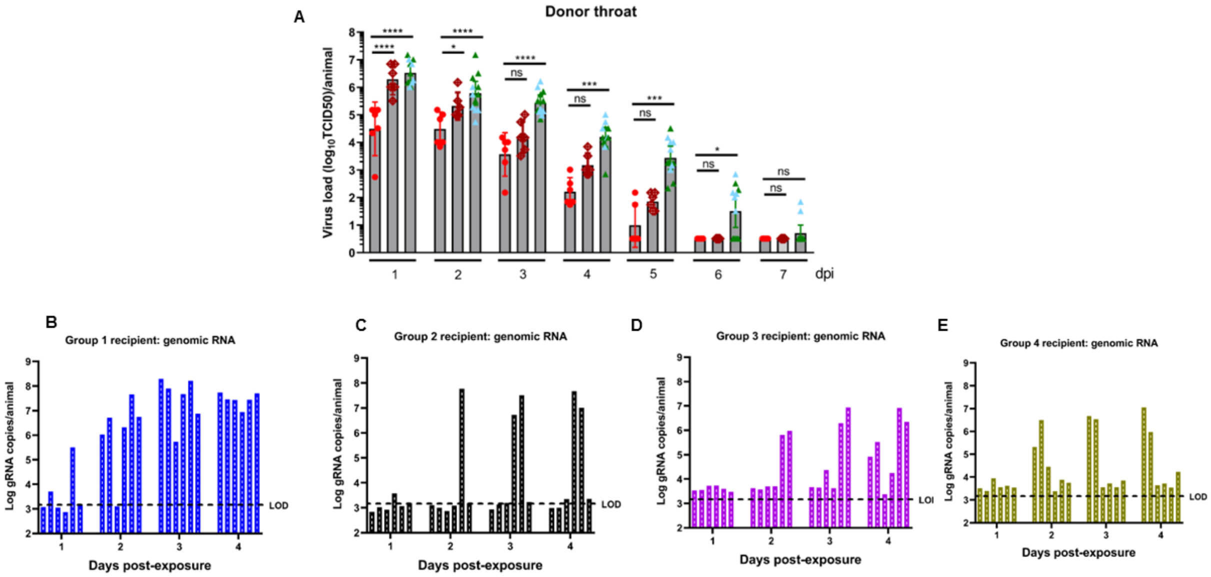


**Extended Data Fig 8. Comparison of virus load and TNA levels in the throat swab**. **A.** Comparison of virus load in the throat swab of the donor hamsters from different groups after the SARS-CoV-2 challenge. The presence of the infectious live virus 1-7 days after infection is determined by $TCID_{50}$ assay in VAT cells. One-way ANOVA followed by Dunnett's multiple comparison test was used for statistical analysis. **B-E**. SARS-CoV-2 RNA levels in throat swabs. Total RNAs were isolated and subjected to the one-step qRT-PCR analysis. Viral loads were quantified as SARS-CoV-2 N gene RNA in throat swab fluid on days 1-4 after exposure. Viral RNA was expressed as N gene RNA copy numbers from each swab or animal, based on an RNA standard included in the assay. LOD, the limit of detection.

Extended Figure 9

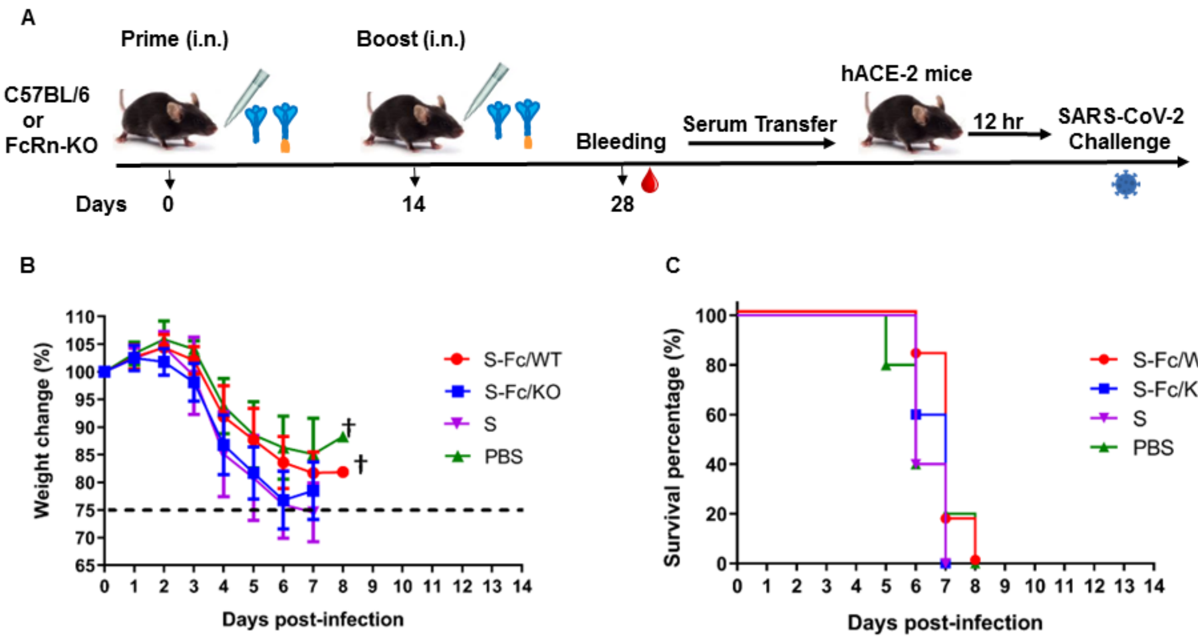


**Extended Data Fig. 9 Evaluation of the passive protection by serum transfer from the immunized mice.** Ten µg of S-Fc, S (with the equivalent molar number), or PBS in combination with 10 µg of CpG was i.n. administered into 6-8-week old wild-type (WT) or FcRn knockout (KO) mice. Mice were boosted 14 days later after primary immunization. Sera were sampled from all immunized mice and 200 µl pooled sera from each group were transferred to 8-week-old hACE-2 mice via i.p injection; 12 hours later, all hACE-2 mice (n=5 or 6 /group) were i.n. challenged with ancestral SARS-CoV-2 (2.5 X $10^4$ $TCID_{50}$) (**A**) and weighed daily for 14 days (**B**). The survival following the virus challenge is plotted as a Kaplan-Meier curve (**C**). Mice were deceased or humanely euthanized if the humane endpoint was reached.

Extended Table 1

**A**

| Recipient | Positive No. after exposure | | | | | | | | | |
|---|---|---|---|---|---|---|---|---|---|---|
| | D1 | D2 | D3 | D4 | D5 | D6 | D7 | D8 | D9 | D10 |
| **Group 1** | 2 | 5 | 6 | 6 | 6 | 6 | 5 | 3 | 0 | 0 |
| **Group 2** | 0 | 1 | 2 | 2 | 2 | 2 | 2 | 1 | 1 | 0 |
| **Group 3** | 0 | 2 | 2 | 4 | 6 | 6 | 6 | 5 | 4 | 4 |
| **Group 4** | 0 | 2 | 2 | 2 | 2 | 2 | 1 | 0 | 0 | 0 |

**B**

<table>
<tr><th rowspan="2">Recipient</th><th rowspan="2">Summarized case No. (P/T)</th><th colspan="4">P values</th></tr>
<tr><th>G1 vs G2</th><th>G1 vs G3</th><th>G1 vs G4</th><th>G2 vs G3</th></tr>
<tr><td>Group 1</td><td>6/6</td><td rowspan="4">0.0303*</td><td rowspan="4">>0.9999</td><td rowspan="4">0.0303*</td><td rowspan="4">0.0303*</td></tr>
<tr><td>Group 2</td><td>2/6</td></tr>
<tr><td>Group 3</td><td>6/6</td></tr>
<tr><td>Group 4</td><td>2/6</td></tr>
</table>

**Extended Table 1. (A)** The infection number of recipient hamsters after exposure to the infected hamsters (donor) was summarized. D: day. **(B)** Statistical differences in the infection rate among different recipient groups were determined by Fisher’s exact test. P: positive number; T: total number. Values marked with asterisks: *, $P < 0.05$.
